# The functional neuronal lipotype is specified by activity and PS synthesis using dietary and lysosomal inputs

**DOI:** 10.64898/2026.09.26.754678

**Authors:** Ryan J. Lee, Irma Magaly Rivas-Serna, Emma K. Theisen, Adriana Bibo, Elizabeth B. Hardy, Peiwen Cheng, Zixiang Zhou, Jocelyn S. Chavez Rios, James C. Forward, Tiffany T. Pham, Connon I. Thomas, Anza Darehshouri, Vera Mazurak, M. Thomas Clandinin, John P. Vaughen

## Abstract

Neurons maintain specialized membranes necessary for lifelong circuit function. While neuronal membranes thus contain unique lipidomes, how and why specific lipids become synaptically enriched remains mysterious. Here, we identify a conserved hallmark of mature neurons, phosphatidylserines containing omega-3 fatty acids (n-3 PS), which dynamically accumulate during synaptogenesis from *Drosophila* to humans. This neuronal lipotype depends on dietary n-3 and lysosomal catabolism mediated by Saposin, the phospholipase PLA2G15, and the lysophospholipid transporter spin/SPNS1. Reducing neuronal PS biosynthesis by targeting the PS synthase (Pss) caused lipidomic, synaptic, and behavioral deficits exacerbated by impaired glycerophospholipid catabolism. While dietary, lysosomal, and lipoprotein manipulations broadly altered n-3 levels, silencing developmental activity-templating neurons selectively reduced n-3 PS. Remarkably, pathogenic *Pss* variants lacking feedback inhibition similarly reduced n-3 PS and impaired circuit function. As n-3 PS is reduced during neurodegeneration but absent from iPSC-derived neurons, important relationships between lipotype and physiology may not be recapitulated *in vitro*.

## Main

The brain contains thousands of diverse lipids which serve biophysical, energetic, and bioactive signaling roles, and which vary across cell-type, organelle, and leaflet^1–3^. For example, Phosphatidylserine (PS) is the dominant negatively charged glycerophospholipid (GPL) that localizes to the inner leaflet of all eukaryotic plasma membranes (Fig. 1a)^4^, and PS externalization can trigger developmental neurite pruning^5,6^. Lipids underlie critical facets of neurobiology, from the fusion and recycling of synaptic vesicle membranes to ion flux through voltage-gated channels encapsulated and tuned by plasma membrane lipid composition^7–9^. Over 50 years ago, brain grey matter, but not white matter, was found to harbor high levels of polyunsaturated fatty acids (PUFA; >1 carbon-carbon double bond)^10,11^, specifically omega-3 (n-3) fatty acids, which are classified by the position of the first double bond from the methyl tail (Fig. 1b, arrow). PUFA chains enable unique membrane properties, including high curvature^12^, fission^13^, and domain stability^14^. Dietary deficiencies in essential PUFAs, or mutations in lipid metabolism enzymes, can cause both neurodevelopmental or neurodegenerative diseases^7,15^. For example, gain-of-function mutations in the PS synthase *PTDSS1* cause a rare disorder, Lenz-Majewski Syndrome^16^, that afflicts bones and skin but also causes intellectual disability, suggesting that regulated PS synthesis is critical for neuronal health. At the same time, our understanding of how lipids are developmentally specified to then sculpt adult nervous system function remains fragmentary.

**Fig. 1:**
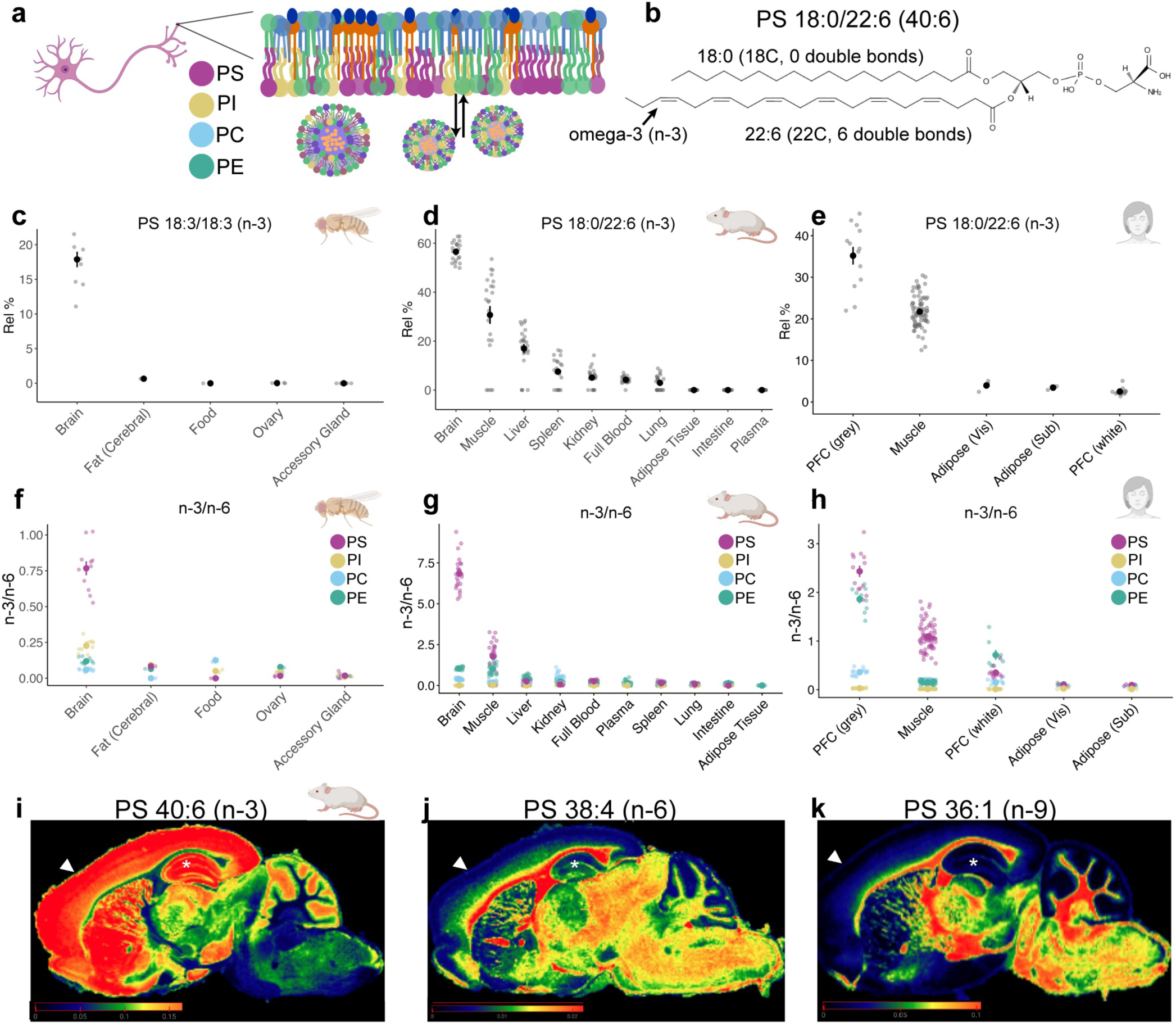
Neurons contain abundant omega-3 phosphatidylserines *in vivo*. **a,** neuronal plasma membrane diagram^1,166^ of phosphatidylserine (PS), phosphatidylinositol (PI), phosphatidylcholine (PC), phosphatidylethanolamine (PE), and sphingolipids (orange). **b,** an omega-3 (n-3) polyunsaturated fatty acid (PUFA) is defined by the position of the first double bond (arrow). **c,** PS 18:3n-3/18:3n-3 relative abundance across fly tissues and diet at day 7. n (tubes) = 10 brains, 3 ovaries, 3 accessory glands, 3 cerebral fat, and 2 control food. **d,** PS 18:0/22:6n-3 relative abundance re-analyzed from mouse tissue^38^ (from P60 mice). n (tubes) = 23 brains; 23 muscle; 23 liver, 24 spleen, 23 kidney, 24 full blood, 24 lung, 24 adipose tissue, 24 intestine, and 24 plasma. **e,** PS 18:0/22:6n-3 relative abundance re-analyzed from human tissues^42,43,167^. n (tubes) = 13 PFC grey; 13 PFC white; 75 muscle; 3 visceral adipose tissue; and 3 subcutaneous fat. **f,** ratio of omega-3 to omega-6 chains (n-3/n-6) in PS, PI, PE, and PC GPL, plotted by tissues in fly **g,** mouse^38^, and **h,** human^41–43^; same n as in c-d. (n-3/n-6 = 18:3/18:2 for fly and (22:6+20:5+18:3)/(20:4+22:4+18:2) for mouse and human). **i,** MALDI-MSI^44^ re-analysis of PS 40:6(n-3); **j,** PS 38:4(n-6); and **k,** PS 36:1(n-9) from a 6 month female mouse control brain (arrows, prefrontal cortex; *, hippocampus). Data are represented by mean and SEM. See Supplemental Table 1 for complete genotypes.

The primary organelle for lipid catabolism is the lysosome, which processes complex membranes containing sphingolipids, sterols, and glycerophospholipids (GPLs) via a battery of lipid-binding proteins and substrate-specific hydrolases^17^. Accordingly, lysosomal storage disorders, in which undigested lipid or protein substrates accumulate in the lysosome, can cause neurodevelopmental disorders and predispose to future neurodegenerative risk^18–20^. Neurodevelopment entails massive membrane rearrangements, from axon guidance to synaptic vesicle biogenesis and myelin wrapping^7^, that are anticipated to require high rates of both lipid biosynthesis and catabolism. While developmental axon pruning and neurogenesis are correlated with altered lysosomal activities^21,22^, how specific outputs of lysosomal lipid catabolism contribute to neurodevelopment and physiology remain incompletely understood. One class of lysosomal proteins implicated in human disease are Saposins (Sphingolipid activator proteins), which are defined by characteristic 4-5 alpha helix domains stabilized by ∼3 disulfide bridges^23^. The best studied human Saposin protein, Prosaposin (PSAP), is synthesized as a proprotein and cleaved in the lysosome into modular effector proteins that facilitate sphingolipid catabolism^24^. PSAP homolog loss in worms, flies, and mice causes lysosomal engorgement and neurological deficits attributed to sphingolipid accumulation^25–30^. While many studies have thus focused on sphingolipids as underpinning disease, Saposins can also bind GPLs *in vitro*^17, 19^, which are abundant yet understudied constituents of neuronal plasma membranes^1,34^ and synaptic vesicles^35^ (Fig. 1a).

Here, we reveal that neurons accumulate uniquely high levels of phosphatidylserines containing n-3 fatty acids (n-3 PS) during development across flies, mice, and humans. In *Drosophila*, the developmental diet is essential for generating the adult brain lipotype: PUFA-free diets imposed in development but not adulthood severely reduce n-3 PS, and developmentally acquired n-3 fatty acids perdure for at least 50 days (>60% of the lifespan). Alongside developmental diet, we identify Saposin-related (Sap-r), the fly homolog of PSAP, as a critical regulator of PUFA accretion into PS (PUFA PS) and other GPLs. Sap-r functions from developing neurons and glia to regulate lysosomal homeostasis and to coordinate PS metabolism. Surprisingly, *Sap-r* phenotypes are not rescued by alleviating sphingolipid burden. Instead, Sap-r functions with homologs of the lysosomal phospholipase PLA2G15 and the lysophospholipid transporter spin/SPNS1 for PUFA PS accumulation in the brain. Broadly reducing PS biosynthesis in developing neurons by targeting the PS synthase *Pss* disrupted the distribution of the synaptic vesicle protein Synapsin and also impaired adult behavior. Strikingly, the dynamic accretion of n-3 PS occurs following the onset of spontaneous and patterned developmental activity consisting of calcium sweeps^36^, and silencing this developmental network activity selectively depleted n-3 PS. Remarkably, inducing pathogenic Lenz-Majewski *Pss* variants into the circuits which template neurodevelopmental activity also selectively reduced n-3 PS levels and impaired circuit function. While n-3 PS and other n-3 GPLs are reduced in frontotemporal dementia patients, n-3 fatty acids are notably absent across multiple neuronal culture models, which instead accumulate lipid hallmarks of essential fatty acid deficiency. Together, these data illuminate how differentiating neurons achieve unique lipid compositions required for synaptic function, with developmental diet and lysosomal inputs enabling lipotype specification via neuronal activity and regulated PS synthesis.

## Results

### Neurons contain abundant omega-3 phosphatidylserines *in vivo*

While profiling lipids in adult *Drosophila* brains using liquid chromatography-mass spectrometry (LC-MS/MS; see Extended Methods), we observed that phosphatidylserines (PS) containing omega-3 (n-3) fatty acids were present at high levels in the brain but not in cerebral adipose or reproductive tissues (Fig. 1c). In *Drosophila*, 18:3 (α-linolenic acid) represents the dominant n-3 fatty acid, while in vertebrates the major n-3 fatty acid is 22:6 (docosahexaenoic acid, DHA)^37^. By analyzing published lipidomics datasets from mice^38–40^ and humans^41–43^, we found that the relative fraction of PS containing 18:0/22:6 was highest in the brain (Fig. 1d-e). Notably, in human prefrontal cortex (PFC), 18:0/22:6 accounted for 35% of PS in grey matter but only 2.5% of PS in white matter (Fig. 1e), suggesting synaptic enrichment of n-3 PS. Indeed, computing the ratio of n-3/n-6 fatty acids across GPLs revealed striking elevations for this ratio in PS (n-3/n-6 PS) compared to other GPLs in the fly brain (Fig. 1f). High n-3/n-6 PS was similarly observed in mouse and human brains, with muscle containing the second highest ratio, and human grey PFC also containing high n-3/n-6 phosphatidylethanolamine (PE; Fig. 1g-h). These ratios largely reflect the contributions of DHA (n-3) and arachidonic acid (AA, 20:4n-6) (Extended Data Fig. 1a-b). In mice, the n-3/n-6 PS ratio was lowest in myelin-rich regions such as the corpus callosum and optic nerve (Extended Data Fig. 1c)^2^. Consistent with stark regional differences between white and grey matter, re-analysis of published imaging MS datasets^44^ revealed high levels of n-3 species such as PS 40:6 (predicted 18:0/22:6) in neuronal regions (Fig. 1i), whereas PS predicted to contain n-6 and n-9 chains were largely restricted to white matter tracts (Fig. 1j-k; Extended Data Fig. 1d-f and^45^). Thus, across species, the brain contains abundant n-3 PS compartmentalized in neurons.

### Neurodevelopmental accumulation of dietary n-3 chains in PS

We next generated a lipidomics dataset from developing *Drosophila* brains to reveal the temporal kinetics of n-3 PS accumulation. Intriguingly, elevated n-3 PS emerged during later brain development, rising from mid-development (48 hours after puparium formation, 48 hAPF) to reach persistently high levels during adulthood (Fig. 2a). Notably, all GPLs dynamically accumulated n-3 and n-6 PUFAs during the last twenty-four hours of neurodevelopment, a period of spontaneous electrical activity and synapse deposition^36,46^ (Extended Data Fig. 2a-g). This developmental PUFA accretion was most pronounced in PS and did not temporally correlate with changes in total PS levels (Extended Data Fig. 2h-j). By mining human^11^ and mice^39^ longitudinal brain lipidomics datasets, we observed a similar accretion of n-3 PS during development that persisted for life (Fig. 2b-c). Across species, this resulted in PS containing the highest n-3/n-6 ratio (Fig. 2d-f), while PI contained the highest relative percent of n-6 chains (Extended Data Fig. 3a-c). In contrast to human grey matter, white matter levels of n-3 PS rapidly declined from fetal stages to adulthood (Extended Data Fig. 3j-l)^11^.

**Fig 2:**
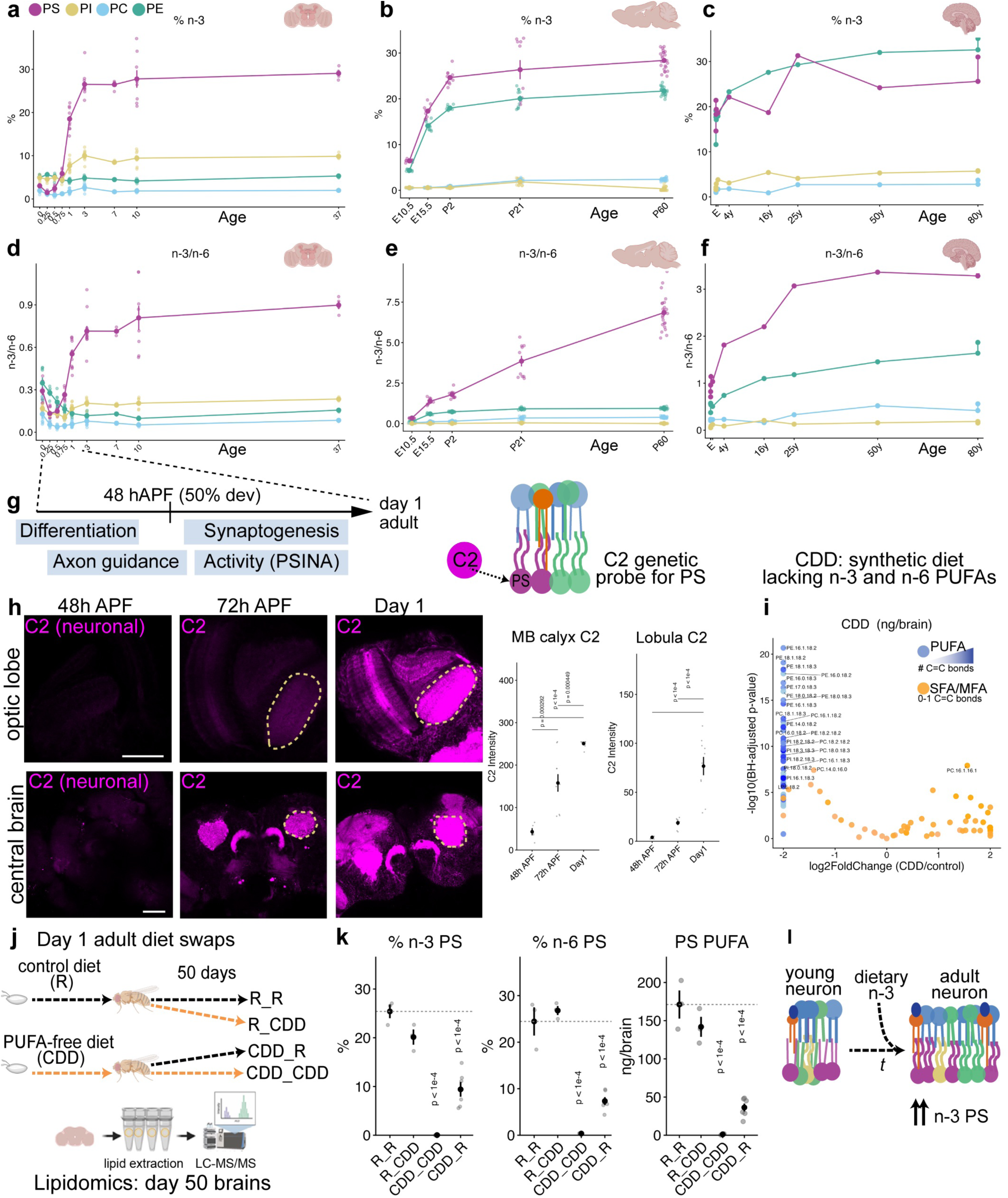
Neurodevelopmental accumulation of dietary n-3 chains in PS. **a,** Glycerophospholipid (GPL) % n-3 distribution in brains dissected across development and adulthood in *Drosophila.* 0 = wandering larval stage L3, 0.25 = 24 hours after puparium formation (hAPF), 0.5 = 48 hAPF, 0.75 = 72 hAPF; 1-37 = day 1 to day 37 adults. n = 8 tubes for all ages except 7 and 37, where n = 4 tubes. **b,** GPL % n-3 distribution across mice brain (cerebral hemisphere) development^38,39^; n = 10 tubes per all ages, except 23 tubes for P60. **c,** % n-3 GPL distribution in human cortex grey matter over age^11^ (n = 1 healthy control brain per listed age; E includes 12 weeks, 35 weeks, 38 weeks, 1 month, and 7 months; other ages indicate years). **d-f,** n-3/n-6 ratios in the brain across age in flies, mice, and humans (same n as a-c). **g,** schematic of *Drosophila* neurodevelopment. **h,** genetically encoded probe for PS (*C2^mScarlet^*^3^, magenta) expressed in neurons (*nSyb-GAL4*) imaged at 48h APF, 72h APF, and day 1 adults, quantified in lobula plate and mushroom body calyx (dashed areas). Scale bar = 50µm. n > 5 brains per timepoint; p-values indicate Tukey’s ANOVA calculated between 48 hAPF and other timepoints. **i,** volcano plots of brain GPLs from day 1 animals reared on CDD (PUFA-free) vs control diets (n=4 tubes/condition); PUFA-containing GPLs are colored blue. 18:2(n-6) decreased from 15.9% to 0.16%, and 18:3(n-3) decreased from 4.8% to 0%. **j,** animals reared on control (R) or PUFA-free (CDD) diets were grown for 10 days into 1 day adults, then maintained on identical or reciprocal diets, before aging for 50 days and profiling brain GPLs. **k,** % n-3 PS, % n-6 PS, and total ng/brain PUFA PS shown for the four dietary conditions; n = 3 tubes for R_R, 3 tubes for R_CDD, 3 tubes for CDD_CDD, and 6 tubes for CDD_R. p-values indicate BH-corrected student’s t-tests (two-sided) compared to R_R controls. **l,** model. Data are represented by mean and SEM. See Supplemental Table 1 for complete genotypes.

To track PS localization *in vivo* across neurodevelopment, we generated a genetically encoded PS biosensor using a minimal PS binding C2 domain from lactadherin (Fig. 2g-h)^4,5^. Neuronally expressed C2 was low at mid-development (48 hAPF) but increased in synaptic regions in the optic lobe and central brain, accumulating from 72 hAPF to adulthood (Fig. 2h). While C2 was present in synaptic neuropils marked by *brp::GFP* (Extended Data Fig. 3n), this synaptic localization was abrogated via neuronal blockade of PS biosynthesis (Extended Data Fig. 3m) or by mutating the lipid binding residues in C2 (C2^AAA^; Extended Data Fig. 3n). Thus, PS localizes to maturing synapses in neurodevelopment.

We next interrogated the dietary underpinnings of n-3 PS accumulation in the brain, as the requisite desaturases for synthesizing 18:3 are absent in flies^47–49^, rats^50,51^, and humans^52–55^. We used either our control cornmeal-molasses diet (R) or a chemically defined diet lacking PUFAs (CDD)^56^, which severely depleted brain n-3 and n-6 fatty acids (Fig. 2i and Extended Data Fig.3o). We raised animals on these diets during development, and then either maintained or swapped diets from day 1 to day 50 of adulthood (∼60% of lifespan) before quantifying brain GPLs (Fig. 2j). As expected, brains from animals solely maintained on CDD diets were devoid of n-3 PS or n-6 PS (Fig. 2k; Extended Data Fig. 3p). Strikingly, animals shifted from control to CDD diets at day 1 of adulthood maintained high levels of PUFA PS throughout life, whereas animals shifted from CDD to control diets failed to accumulate wild-type levels of PUFA PS, even after 50 days (Fig. 2k; Extended Data Fig. 3p). Thus, the developmental diet plays an essential and long-lasting role in specifying the adult neuronal lipotype (Fig. 2l). Notably, given that our standard diet is devoid of the major n-3 PS (Fig. 1c) and that there is no dietary consumption during metamorphosis, a genetic program must redistribute essential n-3 fatty acids into neuronal PS at the correct developmental time.

### *Saposin* mutants reduce PUFA-containing phospholipid levels in the brain

What genes and molecular mechanisms regulate the conserved neuronal lipotype of abundant n-3 PS? While screening brain fatty acids by LC/MS-MS in lipid-metabolic perturbations (including *gba1b, cerk, schlank, lace, bwa, norpA, Asm,* and *iPLA2-VIA*), we unexpectedly found that PUFA GPL chains were extensively reduced in *Sap-r^Δ^*brains (Fig. 3a-c; Extended Data Fig. 4a-b). In contrast, monounsaturated (MUFA) GPLs increased across all classes in *Sap-r^Δ^*brains (Fig. 3b; Extended Data Fig. 4c-h), as did most phosphatidylinositol (PI) species (Extended Data Fig. 4i-j). While mammalian Prosaposin (PSAP) is a lipid chaperone known to potentiate sphingolipid catabolism, including by facilitating glucocerebrosidase activity (GBA; gba1b) (Fig. 3a), *gba1b^Δ^* did not reduce PUFAs in the brain (Fig. 3b). Profiling *Sap-r^Δ^* brain lipids across development and adulthood revealed that PUFA PS species were persistently reduced by ∼38% (n-3 PS) and ∼33% (n-6 PS) from juvenile ages onwards (Fig. 3c), a larger decrease than in other PUFA GPLs (Extended Data Fig. 4d-e). To test if dietary supplementation could bypass Sap-r function, we fed *Sap-r^Δ^* animals diets containing 0.1% or 0.5% flaxseed oil (FO) or walnut oil (WO), which contain abundant 18:3n-3 chains (Fig. 3d). In both control and *Sap-r^Δ^* animals, oil-supplemented diets strongly elevated n-3 PS and other n-3 GPLs in the brain, yet *Sap-r^Δ^* animals consistently accumulated less n-3 PS than controls across most dietary conditions, particularly in 0.1% FO and 0.1% WO regimes (Fig. 3e; Extended Data Fig. k-l); additionally, in all FO diets, n-6 GPLs were strongly depleted. Thus, while *Sap-r^Δ^* can be bypassed by increasing dietary n-3 bioavailability, the maximal extent of dietary n-3 incorporation into the brain appears dependent on Sap-r function.

**Fig. 3:**
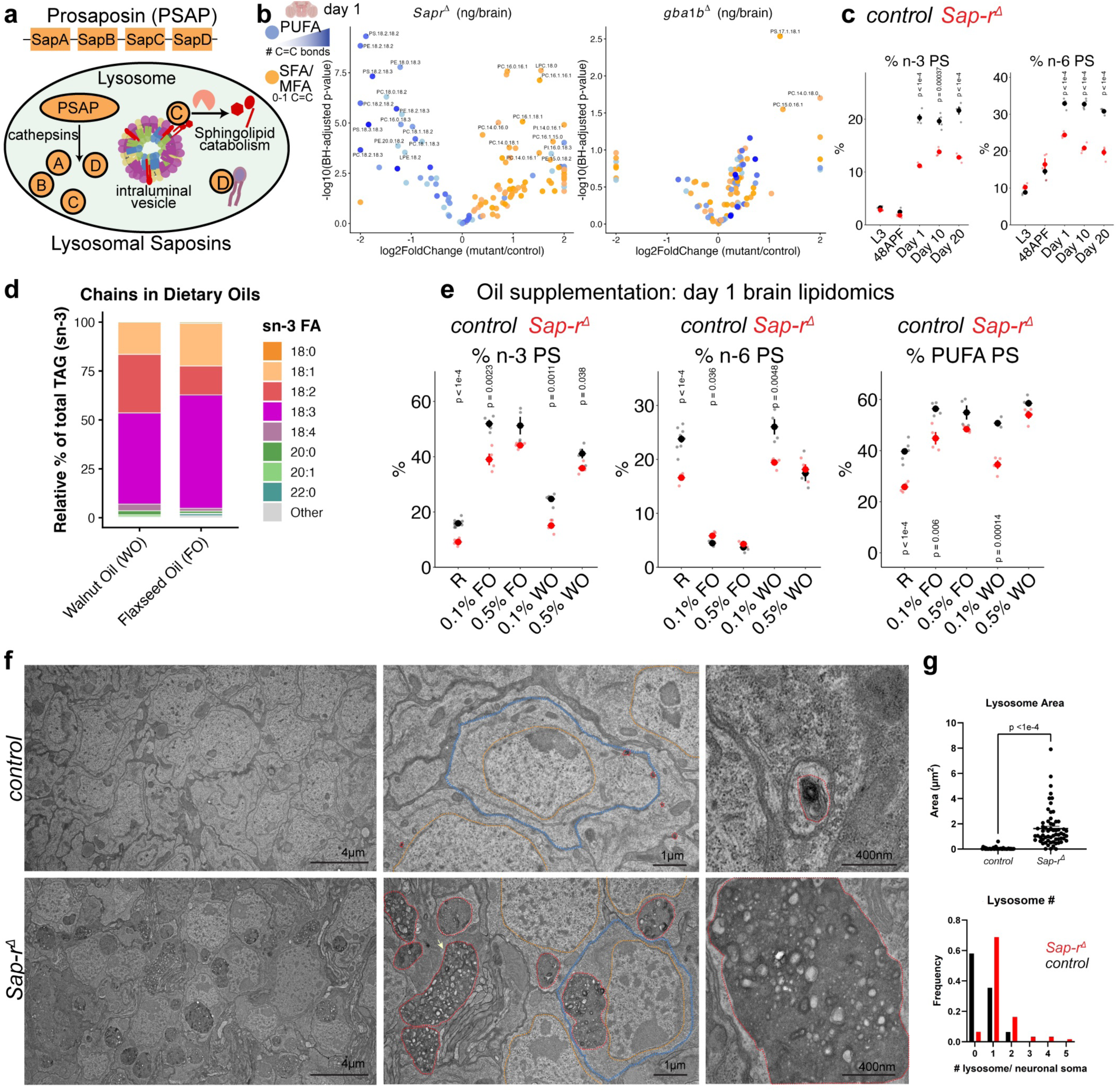
Lysosomal *Saposin* mutants reduce polyunsaturated phospholipids in the brain. **a,** model of human Prosaposin (PSAP) function: lysosomal cathepsins cleave PSAP into modular domains thought to potentiate sphingolipid catabolism by interfacing with intraluminal vesicles. **b,** volcano plot of brain GPL from *Sap-r^Δ22/10^* (hereafter Sap-r*^Δ^*) versus controls at day 1, or *gba1b^ΔTT/Δ^*^1^ versus controls at day 1 (n = 4 tubes per genotype); blue = PUFA-containing GPLs, orange = MUFA and SFA containing GPLs. Annotated species are p<0.05 by BH-adjusted t-test, and >0.5 log2 fold-change. **c**, % n-3 PS and % n-6 PS in *Sap-r^Δ^*vs control brains over development (L3, 48h APF, and Day 1) and into adulthood (Day 10 and Day 20). Note axis age is a categorical and not continuous variable. n = 4 tubes per genotype, p-values indicate two-sided t-test comparison of genotypes at a given age. **d,** dietary oil fatty acid chain composition (sn-3 chains in triacylglycerols) from Walnut Oil (WO) or Flaxseed oil (FO). **e,** control and *Sap-r^Δ^* (red) animals were reared on control (R), 0.1% FO, 0.5% FO, 0.1% WO, or 0.5% WO, and brains subjected to lipidomics at day 1, with % n-3, % n-6, and % PUFA PS graphed (n = 4 tubes/condition; p-values indicate two-sided t-test comparison of genotypes for a given diet). **f,** electron microscopy of day 1 control and *Sap-r^Δ^* optic lobe soma (red lines = lysosomes, orange lines = neuronal nucleus; blue lines = example soma; arrow = glial lysosomes; scale bars = 4µm, 1µm and 400nm). n = 5 brains and 10 sections. **g,** quantification of lysosomal area and frequency. p-value indicates two-sided student’s t-test conducted in PRISM. Data are represented by mean and SEM. See Supplemental Table 1 for complete genotypes.

To observe the subcellular consequences of *Sap-r^Δ^* on the brain, we conducted electron microscopy (EM) of optic lobe neurons in juvenile (day 1) flies. Consistent with Sap-r functioning in lysosomal lipid catabolism, enlarged lysosomes replete with multivesicular and multilamellar structures pervaded *Sap-r^Δ^* mutant soma, with most neuronal soma containing a single engorged lysosome in these 2D slices (Fig. 3f-g). While disrupting sphingolipid catabolism can trigger similar lysosomal defects^57–59^, targeting *gba1b* or *acid sphingomyelinase* did not reduce PUFA PS in the brain (Fig. 3b; Extended Data Fig. 4a). Thus, among the other tested lysosomal mutants, Sap-r appeared uniquely critical for accumulating diet-derived PUFAs into PS during neurodevelopment.

### Sap-r is required in developing neurons and glia for PUFA PS metabolism

To identify if cell types in the brain autonomously required Sap-r, we restricted *Sap-r* knockdown (*Sap-r^KD^*) or rescue manipulations to developing glia (*repo-GAL4*) or developing neurons (*elav-GAL4*); the *RNAi* mediating *Sap-r^KD^* had 70% knockdown efficiency in the brain (Extended Data Fig. 5a). Consistent with EM analyses, LysoTracker+ acidic compartments increased in *Sap-r^Δ^* optic lobe soma (Fig. 4a), and could be significantly but incompletely rescued by overexpressing Sap-r in either neurons or glia (Fig. 4c). Conversely, *Sap-r^KD^* in neurons broadly increased LysoTracker, while glial removal sparsely increased LysoTracker (Fig. 4b-c), perhaps reflecting that glia account for ∼10% of the fly brain^60^. Co-expression of a synthetic *UAS-Sap-r* rescued *Sap-r^KD^* LysoTracker phenotypes (Extended Data Fig. 5b). Lipidomics further revealed that neuronal or glial re-expression of Sap-r rescued PUFA PS in *Sap-r^Δ^* mutants (Fig. 4d and Extended Data Fig. 5c), demonstrating genetic sufficiency for brain-derived Sap-r to mediate PUFA PS accretion. Individually depleting *Sap-r* from neurons or glia weakly reduced PUFA PS, while co-depletion significantly reduced PUFA PS (Fig. 4d) and increased total PI (Extended Data Fig. 5e). Ubiquitous *Sap-r* knockdown also reduced PUFA PS, yet preventing glial knockdown in this background (*repo-GAL80*) partially rescued the PUFA PS deficit (Extended Data Fig. 5e). Thus, while Sap-r exerts some level of cell autonomous control on lysosomal homeostasis and PUFA PS, non-autonomous interactions between developing neurons and glia (and potentially other tissues) may also contribute to the developmental accretion of PUFA PS in the brain. These data implicate Sap-r function in both developing neurons and glia, consistent with its scRNAseq expression^61^ and with endogenous transcriptional GAL4 reporters (Extended Data Fig. 5f, *Sap-r-T2A-GAL4*^62^).

**Fig. 4:**
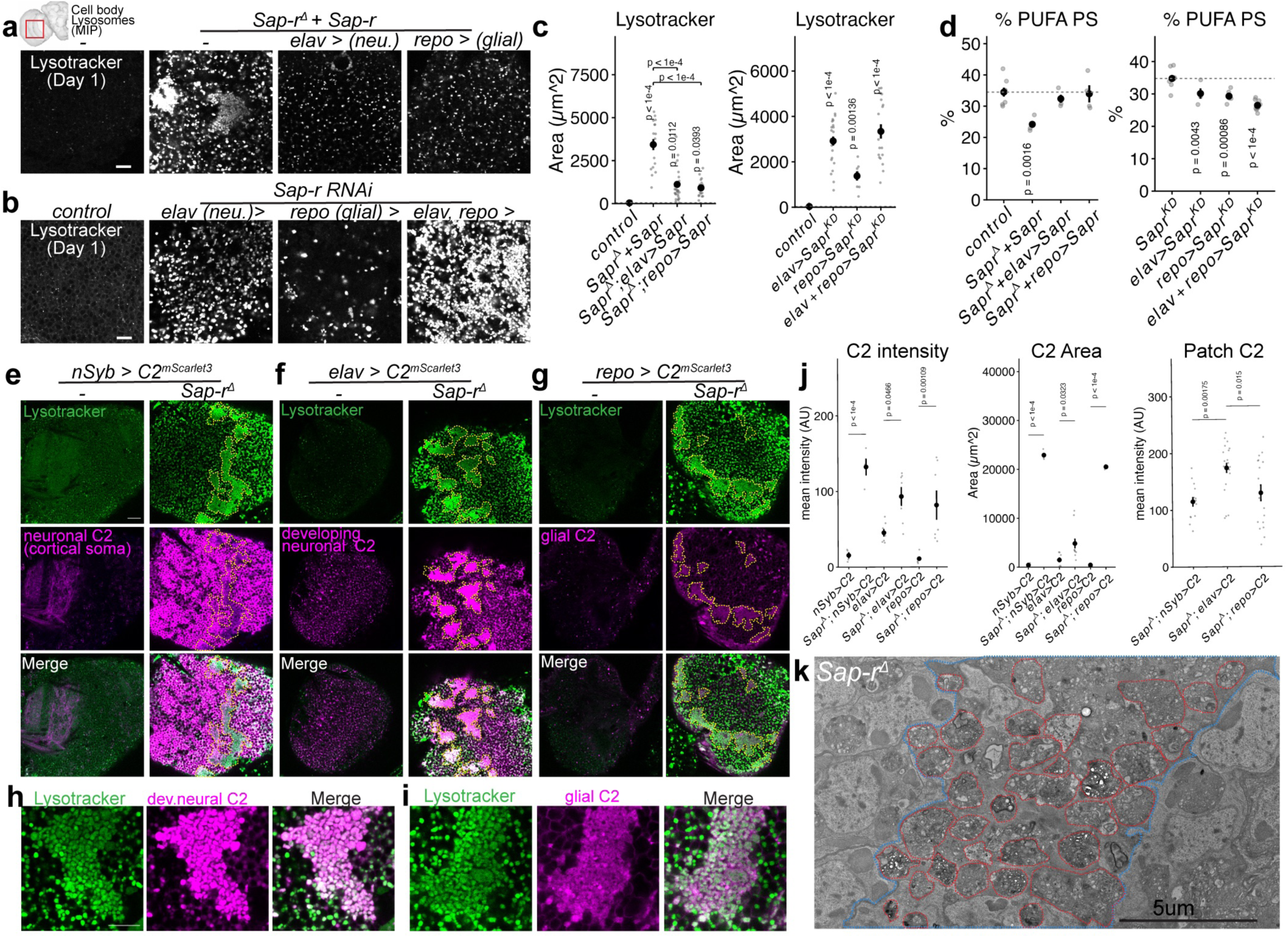
Sap-r is required in developing neurons and glia for PUFA PS metabolism. **a-c,** LysoTracker staining of optic lobe soma upon neuronal (*elav*) or glia (*repo*) *Sap-r* rescue or knockdown. Scale bar = 20µm. Rescue: n (optic lobes) = control (8), *UAS-Sap-r; Sap-r^Δ^* (18); *elav* > *UAS-Sap-r; Sap-r^Δ^* (30) and *repo* > *UAS-Sap-r; Sap-r^Δ^* (18). Knockdown: n (optic lobes) control (19), *elav>Sap-r^KD^* (26), *repo > Sap-r^KD^*(8), *elav, repo > Sap-r^KD^*(19). p-values indicate BH-corrected pairwise t-tests (two-sided) between controls and genetic perturbations; mean and SEM are shown. **d,** brain PUFA PS % in cell type-specific *Sap-r* rescue or knockdowns. Lipidomics rescue n (tubes): 8 control tubes, and 4 tubes for all other genotypes. Lipidomics knockdown n (tubes): control (8); *elav* (4); *repo* (5); and *elav, repo* (10). p-values indicate BH-corrected pairwise student’s t-tests (two-sided) between controls and genetic perturbations; mean and SEM are shown. **e-i,** day 1 control and *Sap-r^Δ^* expressing *C2^mScarlet^*^3^ (magenta) in developing (*elav*) or maturing (*nSyb*) neurons or glia (*repo*), stained with LysoTracker (green). Scale bar = 20µm. **j,** intensity quantification of C2 mean intensity and area per optic lobe, and mean intensity in LysoTracker patches. n (optic lobes) = 4 *nSyb>C2*; 4 *nSyb>C2; Sap-r^Δ^* ; 4 *repo>C2*; 3 *repo>C2; Sap-r^Δ^* ; 7 *elav>C2*; and 11 *elav>C2; Sap-r^Δ^*. p-values from two-sided t-tests between indicated control and *Sap-r* conditions. **k,** electron microscopy of putative LysoTracker patches (blue boundary) in *Sap-r^Δ^* optic lobe at day 1; subsets of lysosomes are outlined in red. n = 5 brains and 10 sections for EM. Scale bar = 5 µm. Data are represented by mean and SEM. See Supplemental Table 1 for complete genotypes.

To track PS in controls and *Sap-r^Δ^*, we expressed the C2 probe in developing neurons (*elav*-*GAL4*), developing glia (*repo*-*GAL4*), or maturing neurons (*nSyb*-*GAL4*), and dissected brains at day 1. Across all cell types, C2 accumulated in distinct patterns across *Sap-r^Δ^*optic lobe cell bodies (Fig. 4e-g; note that the majority of neuronal C2 is synaptic, Fig. 2h). Strikingly, *elav*-derived C2 formed smaller puncta in discrete patches that colocalized with LysoTracker (Fig. 4h; “LysoTracker patches”). LysoTracker patches were largely devoid of *nSyb*-derived C2 (Fig. 4e) but colocalized with glial C2 (Fig. 4i), which appeared as an amorphous cytosolic signal enclosing the puncta Lysotracker signal (Fig. 4g-i). We observed similar C2-LysoTracker patterns upon *Sap-r^KD^* in both developing neurons and glia (Extended Data Fig. 5g-i). Using EM to examine the optic lobes of *Sap-r^Δ^* brains in more detail, we identified broad areas of membranous inclusions spanning >150µm^2^, packed densely with clustered lysosome-like organelles containing heterogeneous material, including vesicles, multilamellar bodies, and white inclusions consistent with axonal fragments^63^ (Fig. 4k, red outlines). Based on these observations and the known engulfment of neuronal membranes during neurodevelopment^63–65^, we hypothesized that LysoTracker patches contain developing neuronal lysosomal lipids (including PS) that have been engulfed by glia but are indigestible without Sap-r. Consistent with this model, LysoTracker patches occurred only upon the simultaneous depletion of *Sap-r* in both developing neurons and glia, but not individual cell-type depletions (Extended Data Fig. 5k). Similarly, overexpressing Sap-r in developing neurons (*elav*+) or glia rescued LysoTracker patches (Fig. 4a), whereas overexpressing Sap-r in maturing neurons (*nSyb*+) did not (Extended Data Fig. 5k). Thus, while Sap-r is autonomously required for neuronal and glial lysosomal homeostasis, non-autonomous interactions between developing neurons and glia regulate PS distribution and PUFA PS accretion.

### GPL deficits appear separable from sphingolipid accumulation in *Sap-r*

In prior models of *Saposin* deficiency, sphingolipid accumulation was proposed to precede secondary defects in lipid metabolism (Extended Data Fig. 6a). In this framework, GPL defects present in *Sap-r* mutants would be dependent on sphingolipid accumulation. To test this, sphingolipid levels were lowered in *Sap-r^Δ^* by blunting *de novo* sphingolipid biosynthesis through targeting the homolog of SPTLC2B, *lace*^66^ (Extended Data Fig. 6b). Surprisingly, depleting *lace* in developing neurons or glia did not rescue the enlarged LysoTracker phenotype in *Sap-r^Δ^,* suggesting the LysoTracker phenotype is not sensitive to *de novo* sphingolipid biosynthesis (Extended Data Fig. 6c-d). Moreover, while glial depletion of *lace* reduced the most abundant sphingolipid accumulating in *Sap-r^Δ^* brains^67^, ceramide phosphoethanolamine (CPE), this manipulation failed to rescue PUFA PS levels (Extended Data Fig. 6e-f). Separately, expressing a C-terminal V5 tagged Sap-r in *Sap-r^Δ^* neurons or glia failed to rescue both LysoTracker and PUFA PS, yet ameliorated sphingolipid accumulation (Extended Data Fig. 6g-j). While *lace* or *Sap-r^V^*^5^ manipulations did not perfectly normalize all sphingolipids (Extended Data Fig. 6k-l), these residual sphingolipid defects are significantly less severe than in *glucocerebrosidase* or *acid sphingomyelinase* mutants^58,67^, which also accumulate LysoTracker yet do not reduce PUFA PS levels (Fig. 3b; ^67^). While we cannot exclude the possibilities that sphingolipids undetected by our targeted analyses are the underlying drivers of GPL and lysosomal pathology, or that sphingolipid defects in peripheral tissues may trigger downstream GPL defects in the brain, these data suggest that reduced PUFA PS levels are genetically separable from sphingolipid accumulation in *Sap-r^Δ^* brains.

### Lysosomal catabolism via PLA2G15 and SPNS1 homologs regulates brain PUFA levels

We next considered a model where Sap-r more directly regulates GPL chain composition in the brain via other lysosomal GPL handling proteins. To identify potential candidates, we conducted a two-part *RNAi* screen (Fig. 5a). As proteins acting in a similar pathway to Sap-r should cause lysosomal defects, we first screened LysoTracker in day 1 juvenile brains. Then, in a secondary screen, the strongest hits were analyzed by LC-MS/MS. From this targeted screen (Fig. 5b), we identified *spinster*, the homolog of the lysophospholipid transporter SPNS1^68–70^, and *CG18858/CG31683,* homologs of the acid phospholipase PLA2G15. CG18858 and CG31683 are adjacent duplications (99.5% similarity), with both genes containing the catalytic serine required for the indiscriminate catabolism of lysosomal GPLs^71–73^ (Extended Data Fig. 7a). For brevity, the dual targeting of CG18858 (*PLA2G15a*) and CG31683 (*PLA2G15b*) is referred to as *PLA2G15^KD^* or *PLA2G15^Δ^*. Depleting *spin* or *PLA2G15* in both developing neurons and glia caused strong LysoTracker phenotypes, as did independent mutant alleles (Fig. 5c).

**Fig. 5:**
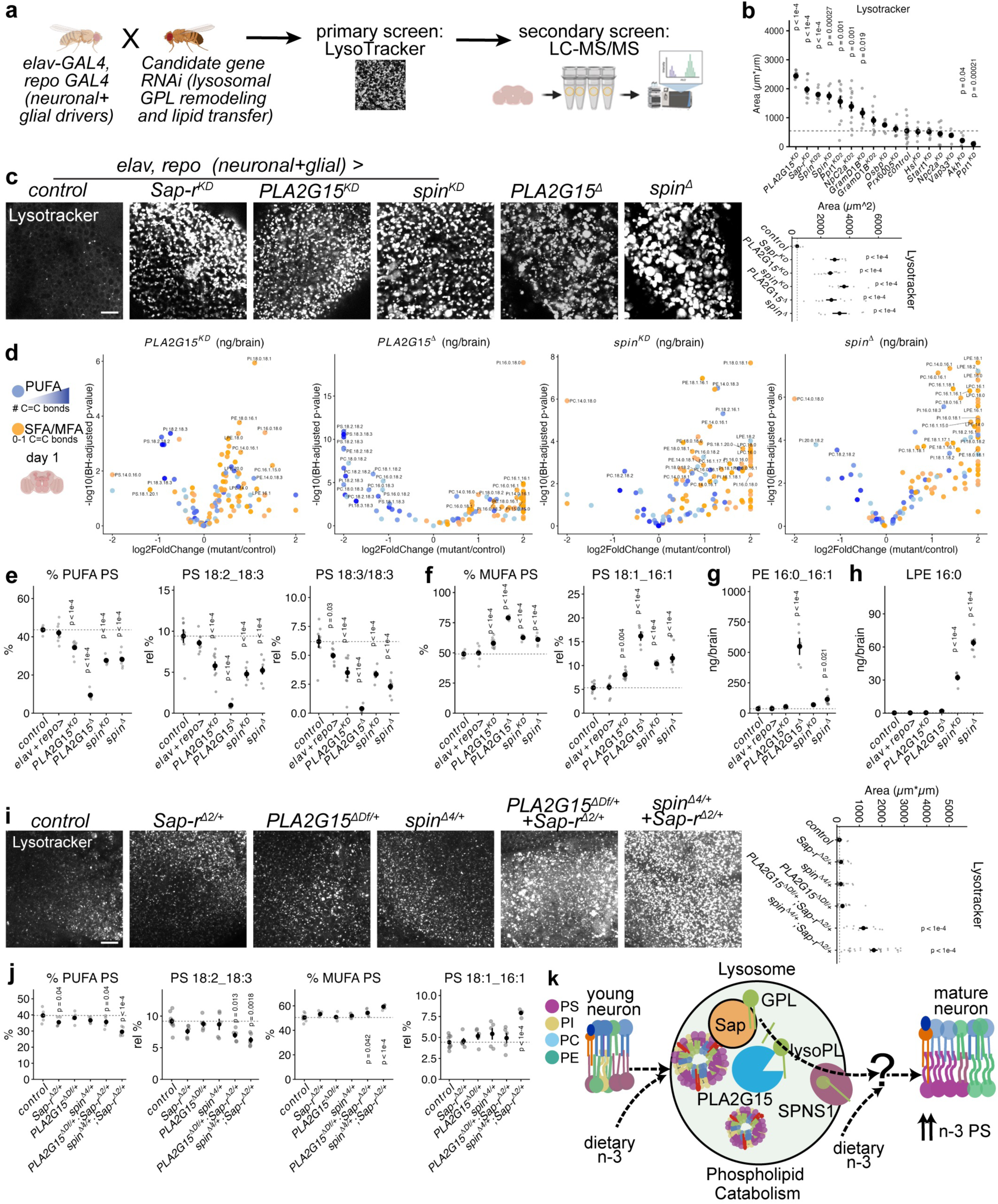
Lysosomal catabolism via PLA2G15 and SPNS1 homologs regulates brain PUFA levels. **a,** screen schematic. **b,** LysoTracker area in optic lobe cortex from a genetic screen using *elav-GAL4,repo-GAL4* in day 1 female brains to drive candidate *RNAi* knockdowns (n ≥ 6 brains per genotype). **c,** day 1 optic lobe soma LysoTracker from controls and screen hits *PLA2G15* and *spin,* mutants and knockdowns. Scale bar = 20µm. n (optic lobes) = 8 control; 7 *Sap-r^KD^*; 12 *PLA2G15^KD^*; 8 *spin^KD^*; 11 *PLA2G15^Δ^*; and 7 *spin^Δ^*. p-values represent BH-corrected two-sided t-tests against control. **d,** volcano plots of phospholipid changes from day 1 brains of indicated genotypes; blue = PUFA, orange = SFA and MFA. For lipidomics, n (tubes) = 8 wild-type, 8 GAL4 control (*elav-GAL4,repo-GAL4*/+); 9 *PLA2G15^KD^*; 4 *spin^KD^*; 5 *PLA2G15^Δ^*; and 5 *spin^Δ^*. For volcano plots, controls were pooled and compared to mutant genotypes with BH-corrected two-sided student t-tests. **e,** major PUFA-containing PS species from screen hits. **f,** major MUFA-containing PS species from screen hits. **g,** levels of PE 16:0_16:1 (ng/brain) and **h,** LPE 16:0 (ng/brain) from screen hits. p-values represent BH-corrected two-sided student’s t-tests against control. **i,** optic lobe LysoTracker staining in transheterozygote combinations of *Sap-r^Δ/+^*with *spin^Δ/+^* or *PLA2G15^Δ/+^*. Scale bar = 20µm. LysoTracker n (optic lobes): 16 control; 17 *Sap ^Δ/+^*; 31 *spin^Δ/+^*; 20 *PLA2G15 ^Δ/+^*; 32 *Sap-r^Δ/+^+spin^Δ/+^* ; 10 *Sap-r^Δ/+^+PLA2G15^Δ/+^*. p-values represent BH-corrected two-sided student’s t-tests against control (dashed line). **j**, lipidomics from day 1 female brains for indicated control and transheterozygous genotypes (n = 5 tubes/mutant genotype, and 8 control tubes). p-values represent BH-corrected two-sided student’s t-tests against control. **k,** model for lysosomal GPL catabolism enabling n-3 incorporation into maturing neuronal PS, with Sap-r facilitating intraluminal lipid transfer, PLA2G15 catabolizing GPLs into lysoPL, and SPNS1 exporting lysoPLs. Data are represented by mean and SEM. See Supplemental Table 1 for complete genotypes.

Paralleling *Sap-r* loss of function, disrupting *spin* or *PLA2G15* broadly lowered PUFA levels in day 1 brains (Fig. 5d; Extended Data Fig. 7b), with significant reductions in major PUFA PS species and increases in MUFA PS (Fig. 5d-f; Extended Data Fig. 7c). Strikingly, *PLA2G15^Δ^* caused the strongest acyl chain defects, with severe depletions of PUFAs and massive accumulation of shorter chain SFAs and MUFAs, leading to large increases in PE and phosphatidylcholine, the most abundant GPLs (Fig. 5g; Extended Data Fig. 5c-d); while *PLA2G15^KD^* did not similarly elevate PE, incomplete knockdown or effects from other tissues may underlie this discrepancy. Consistent with its known biochemical function^68–70^, *spin* perturbations dramatically elevated lysoPEs bearing similar chains to accumulated GPLs in *PLA2G15^Δ^*(Fig. 5h; Extended Data Fig. 7d). Notably, although *Sap-r ^Δ^* mutants are adult viable, adult *spin^Δ^* animals were sub-viable with prevalent pharate lethality (<10% eclosed homozygous mutant), whereas *PLA2G15^Δ^*animals were 100% pharate lethal. Adult *PLA2G15^Δ^* lethality was rescued by global expression of a *PLA2G15a* transgene (Extended Data Fig. 7e), and *PLA2G15* transgenes also rescued lysotracker and lipidomic defects when induced ubiquitously or in the brain specifically (Extended Data Fig. 7f-g). Thus, *spin* and *PLA2G15* loss compromises adult viability, perhaps due to accumulation of lysoPLs and GPLs, respectively. While pronounced PE accumulation in *spin* and *PLA2G15* perturbations are consistent with a model of n-3 fatty acid chains being redistributed from PE to PS during brain development, the total levels of adult brain n-3 PS exceeded those of developmental n-3 PE, implicating additional sources for n-3 chains (Extended Data Fig. 7h).

We next tested for genetic interactions between Sap-r with Spin or PLA2G15 by comparing LysoTracker between heterozygous *Sap-r* mutants and transheterozygous mutant combinations. Strikingly, *PLA2G15^Δ/+^; Sap-r ^Δ/+^* and *spin ^Δ/+^; Sap-r ^Δ/+^* increased LysoTracker in juvenile brains compared to controls and individual heterozygote alleles, with stronger phenotypes in *spin ^Δ/+^; Sap-r ^Δ/+^* animals (Fig. 5i). Lipidomics of transheterozygous brains revealed significant depletion of PUFA PS and other PUFA GPLs and increased MUFA GPLs in *spin ^Δ/+^; Sap-r ^Δ/+^*combinations, with weaker effects detected in *PLA2G15^Δ/+^; Sap-r ^Δ/+^* (Fig. 5j; Extended Data Fig. 7j). In a separate genetic interaction, in a weaker neuronal *PLA2G15* knockdown, LysoTracker intensity was enhanced by *Sap-r ^Δ/+^* (Extended Data Fig. 7i). These data suggest that Sap-r, PLA2G15, and spin may function in a similar genetic pathway, with stronger lipidomic interactions detected between Sap-r and spin. In the simplest model, Sap-r/PSAP facilitates the redistribution of intraluminal GPLs in the lysosome, enabling PLA2G15 catabolism of GPLs to lysoPLs for export via spin/SPNS1 (Fig. 5k), with similar chains accumulating in *PLA2G15 ^Δ^* and *spin ^Δ^* from sequential lipid processing (Fig. 5g-h). In more complicated models, lysosomal functions of these proteins could be spatiotemporally uncoupled in the brain. To further explore the relationships between these lysosomal proteins, we analyzed developing optic lobe scRNA-seq data^61^, which revealed higher glial than neuronal expression for both *Sap-r* and *spin*, with stable neuronal expression throughout development (Extended Data fig. 7k). While *PLA2G15a* and *PLA2G15b* were also expressed in developing glia and neurons, these transcripts dynamically increased during synaptogenesis from 60h APF to peak at 84 hAPF (Extended Data Fig. 7k). Intriguingly, although *PLA2G15a* colocalized well with lysosomal *Sap-r* when induced in neurons (Extended Data Fig. 7l-m), *PLA2G15a* also localized to subsets of brain synaptic neuropil (Extended Data Fig. 7m).

Together, these data reveal that lysosomal Sap-r, PLA2G15, and spin are crucial for generating PUFA PS in the adult brain, and can function from developing neurons and glia.

### Neuronal PS biosynthesis is crucial for brain n-3 PS, Synapsin, and behavior

How do lipids liberated from the lysosome lead to the neuronal accumulation of n-3 PS in the adult brain? LysoPLs are thought to be rapidly re-esterified to GPLs, with select GPLs undergoing conversion to PS in the endoplasmic reticulum via Phosphatidylserine Synthases. In *Drosophila,* the sole PS synthase, Pss, converts PE to PS^74^, and hypomorphic *Pss* alleles cause neurodegeneration^75^. We thus investigated the consequences of selectively depleting *Pss* in developing neurons. Removing *Pss* in neurons (*elav > Pss^KD^*) broadly reduced PS lipids in the adult brain (Fig. 6a). In these animals, we also observed reductions in the area covered by the major synaptic vesicle protein Synapsin^35^ in the mushroom body calyx compared to controls (Fig. 6b); *Pss* hypomorphs also reduced Synapsin area (Extended Data Fig. 8a-b). While strong glial knockdown of *Pss* causes lethality, moderate knockdown regimes that permit adult viability and cause neurodegeneration^75^ did not reduce Synapsin area in the mushroom body (Extended Data Fig. 8a-b), and neither glial nor neuronal *Pss^KD^* altered LysoTracker (Extended Data Fig. 8d). Alongside altering Synapsin coverage, neuronal *Pss^KD^* was immediately deleterious to behavior, with slower climbing times in negative geotaxis assays (Fig. 6c).

**Fig. 6:**
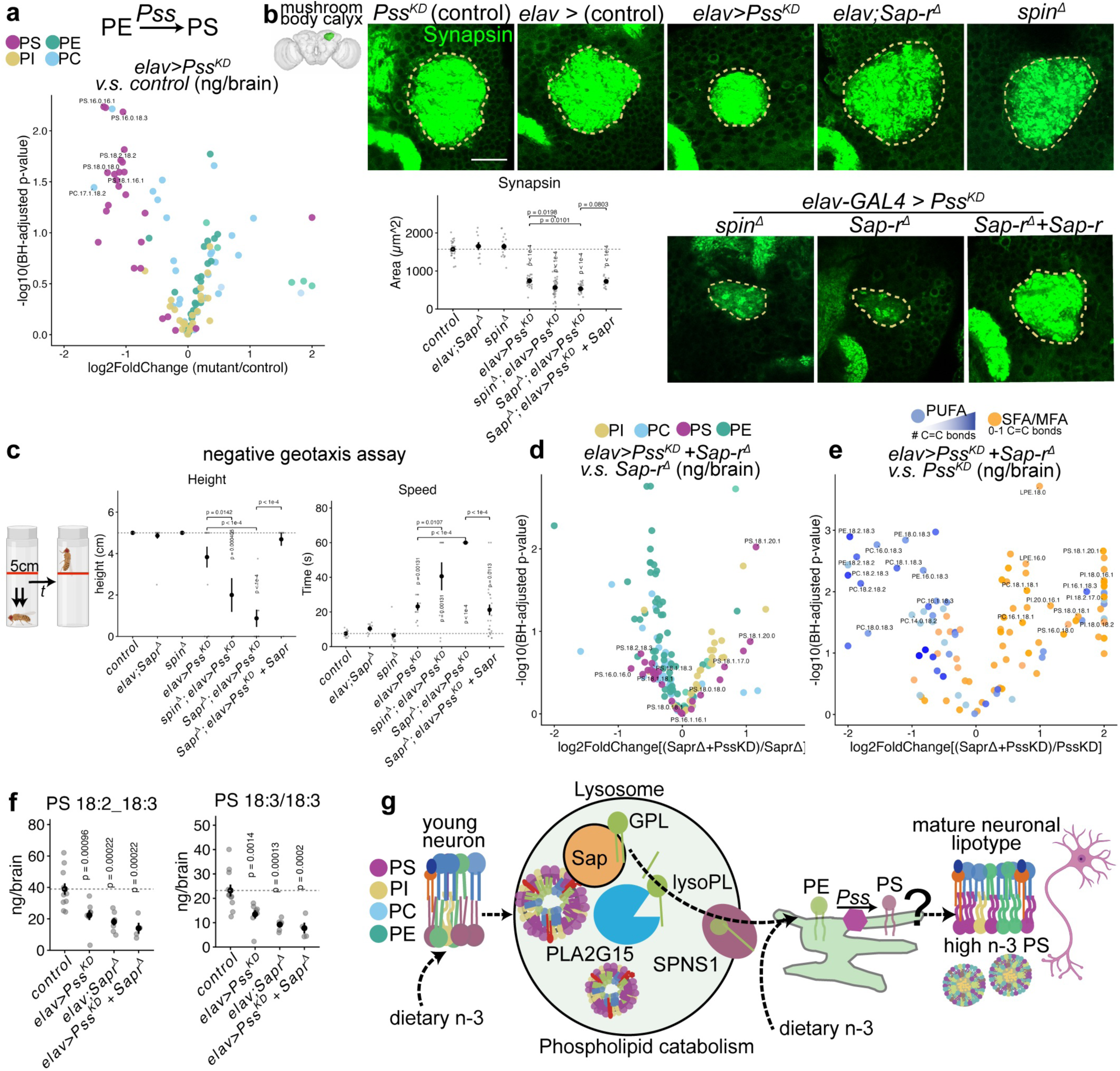
Neuronal PS biosynthesis is crucial for brain n-3 PS, Synapsin, and behavior. **a,** volcano plots of brain GPLs between pooled controls (*elav-GAL4/+* (5 tubes) and *Pss^KD^/+* (6 tubes)) versus *elav>Pss^KD^*(8 tubes), colored by GPL class. **b,** day 1 mushroom body calyx (diagram modified from^168^) stained with Synapsin (green) across indicated genotypes. Scale bar = 20µm. n = 26 control calyxes (*Pss^KD^/+* brains); 15 *spin^Δ^*calyxes; 10 *elav;Sap-r^Δ^* calyxes, 30 *elav >Pss^KD^*calyxes; 37 *elav>Pss^KD^*+ *spin^Δ^* calyxes; 29 *elav>Pss^KD^*+ *Sap-r^Δ^*calyxes; and 17 *elav>Pss^KD^*+ *Sap-r^Δ^* +*Sap-r* brains. p-values represent BH-corrected two-sided t-tests against control. **c,** negative geotaxis assay (day 2 females used). Time and average distance climbed in 60s (capped at 5cm) is quantified. n (flies) = 14 *Pss^KD^/+*; 14 *elav;Sap-r^Δ^*; 15 *spin^Δ^*; 10 *elav>Pss^KD^*+ *spin^Δ^*; 13 *elav >Pss^KD^*; 13 *elav>Pss^KD^*+ *Sap-r^Δ^*; and 26 *elav>Pss^KD^*+ *Sap-r^Δ^* +*Sap-r*. p-values represent BH-corrected two-sided t-tests against control. **d**. volcano plot comparing double mutant *Sap-r^Δ^+elav>Pss^KD^* (4 tubes) to *elav>Pss^KD^* (8 tubes), colored by GPL class; **e,** volcano plot comparing double mutant *Sap-r^Δ^+elav>Pss^KD^* (4 tubes) to *elav;Sap-r^Δ^*(7 tubes), colored by double bond number. **f,** ng/brain of major n-3 PS species in genotypes from f-g. **g,** model: diet and lysosomal catabolism furnish essential lipid substrates for the neuronal lipotype. In the ER, Pss synthesizes PS. Data are represented by mean and SEM. See Supplemental Table 1 for complete genotypes.

We next tested if neuronal *Pss* blockade was sensitive to lysosomal perturbations. While mutating *Sap-r* or *spin* alone did not cause obvious Synapsin defects in the mushroom body calyx, combining *Sap-r^Δ^* or *spin^Δ^*with neuronal depletion of *Pss* enhanced Synapsin loss (Fig. 6b). Strikingly, neuronal *Pss^KD^* climbing defects were potently enhanced by simultaneously mutating *spin* or *Sap-r* (Fig. 6c). Neuronal re-expression of Sap-r in *Sap-r^Δ^+elav >Pss^KD^* ameliorated Synapsin and behavioral phenotypes to the baseline deficit exhibited by *elav >Pss^KD^* (Fig. 6b-c). In contrast, mutating *gba1b* in *elav > Pss^KD^* animals did not impair Synapsin coverage or climbing behavior (Extended Data Fig. 8a-c). Thus, lysosomal GPL catabolism appears to cooperate with neuronal PS biosynthesis, which is essential for Synapsin area and adult behavior. We note that although CDD-reared animals had climbing defects, there were no obvious reductions in Synapsin area (Extended Data Fig. 8a-b). CDD animals also exhibited a severe hypoactivity rescuable by flaxseed oil, with a similar, albeit weaker, phenotype occurring in *Sap-r ^Δ^* animals (Extended Data Fig. 8g-j). Interestingly, while *PLA2G15^Δ^* was pharate lethal and precluded climbing assays, ubiquitous *PLA2G15^KD^*permitted rare “escaper” adults (< 10% eclosion rate) which were immediately defective in negative geotaxis (Extended Data Fig. 8c-d). While *PLA2G15^Δ^* or *PLA2G15^KD^* did not reduce Synapsin area in the mushroom body calyx, the intensities of Synapsin and the active zone protein brp were globally reduced (Extended Data Fig. 8a-e), revealing that loss of *PLA2G15* has severe consequence for synapse development. Separately, we also observed that optic lobe wiring was defective in 100% of *spin^Δ^* adults but not in other lysosomal mutants (Extended Data Fig. 8e, arrows). This may reflect functions for spin in glial migration and subsequent neuropil positioning^76^, and glial-specific *spin^KD^* indeed caused severe lysosomal enlargement (Extended Data Fig. 8f). Thus, although *spin*, *PLA2G15,* and *Sap-r* are essential for lysosomal homeostasis and n-3 PS accumulation in neurodevelopment, differential phenotypes may arise depending on the stage at which lipid catabolism is blockaded.

We also profiled the brain lipidomes of the behaviorally deficient *Sap-r^Δ^*+ *elav > Pss^KD^* animals. Compared to mutating *Sap-r^Δ^* alone, depleting neuronal *Pss* in *Sap-r^Δ^* mutants mildly reduced both PE and PS species (Fig. 6d). Compared to depleting *Pss^KD^* alone, mutating *Sap-r^Δ^* in *Pss^KD^* caused strong reductions in PUFAs, particularly PE and PC species (Fig. 6e). Thus, the effect of neuronal *Pss* depletion appears to be relatively selective for PS, while Sap-r is critical for enabling global PUFA accretion. Whereas individually the *Sap-r^Δ^* or *Pss^KD^* perturbations lowered major n-3 and n-6 PS species, combing these perturbations caused the lowest levels of PUFA PS and other PUFA GPLs (Fig. 6f and Extended Data Fig. 8k-l). Together, these data argue that neuronal PS biosynthesis via Pss, supported by lysosomal GPL catabolism, is indispensable for producing the n-3 PS neuronal lipotype, Synapsin coverage, and adult behavior (Fig. 6g).

### Neuronal activity and Pss feedback inhibition specify the functional neuronal lipotype

As n-3 PS accumulated concordant with developmental neuronal calcium signaling (Fig. 7a), we wondered if developmental neuronal activity instructed lipidomic remodeling. To this end, we focused on the < 2% of neurons that template brain-wide neuronal calcium activity and are molecularly defined by *Trpγ* expression^77^ (Fig. 7b). These *Trpγ+* neurons accumulate the C2 PS sensor throughout neurodevelopment (Extended Data Fig. 9a), and silencing *Trpγ+* neurons via *Kir2.1* severely decreases calcium activity^77^ and causes pharate lethality (Fig. 7b). Lipidomics of *Trpγ>Kir2.1* revealed decreased levels of major n-3 PS species, including PS 18:3/18:3 (Fig. 7c). This lipidomic effect was remarkably selective, as n-6 PS levels were not reduced, nor were other n-3 GPLs reduced (Fig. 7d, Extended Data Fig. 9b). We further tested *Trpγ^Δ^* mutants, which lack the nonselective cation *Trpγ* channel and have altered developmental calcium activity, with blunted calcium amplitude but increased calcium sweeps, and which readily eclose^77^. Surprisingly, *Trpγ^Δ^* mutants harbored increased essential fatty acid chains across all GPLs (Extended Data Fig. 9d), perhaps due to systemic effects on lipogenesis^78^ or to the doubling of calcium sweeps during active phases^77^. Further pointing to complex relationships between calcium and GPL composition, hypomorphic mutants in the voltage-gated calcium channel Cacophony^79^ non-selectively lowered all PUFA GPLs (Extended Data Fig. 9d). Thus, while alterations to calcium homeostasis may broadly tune PUFA levels, the activity pattern of the Trpγ+ neuronal population is essential for the selective accumulation of n-3 PS in the brain.

**Fig. 7:**
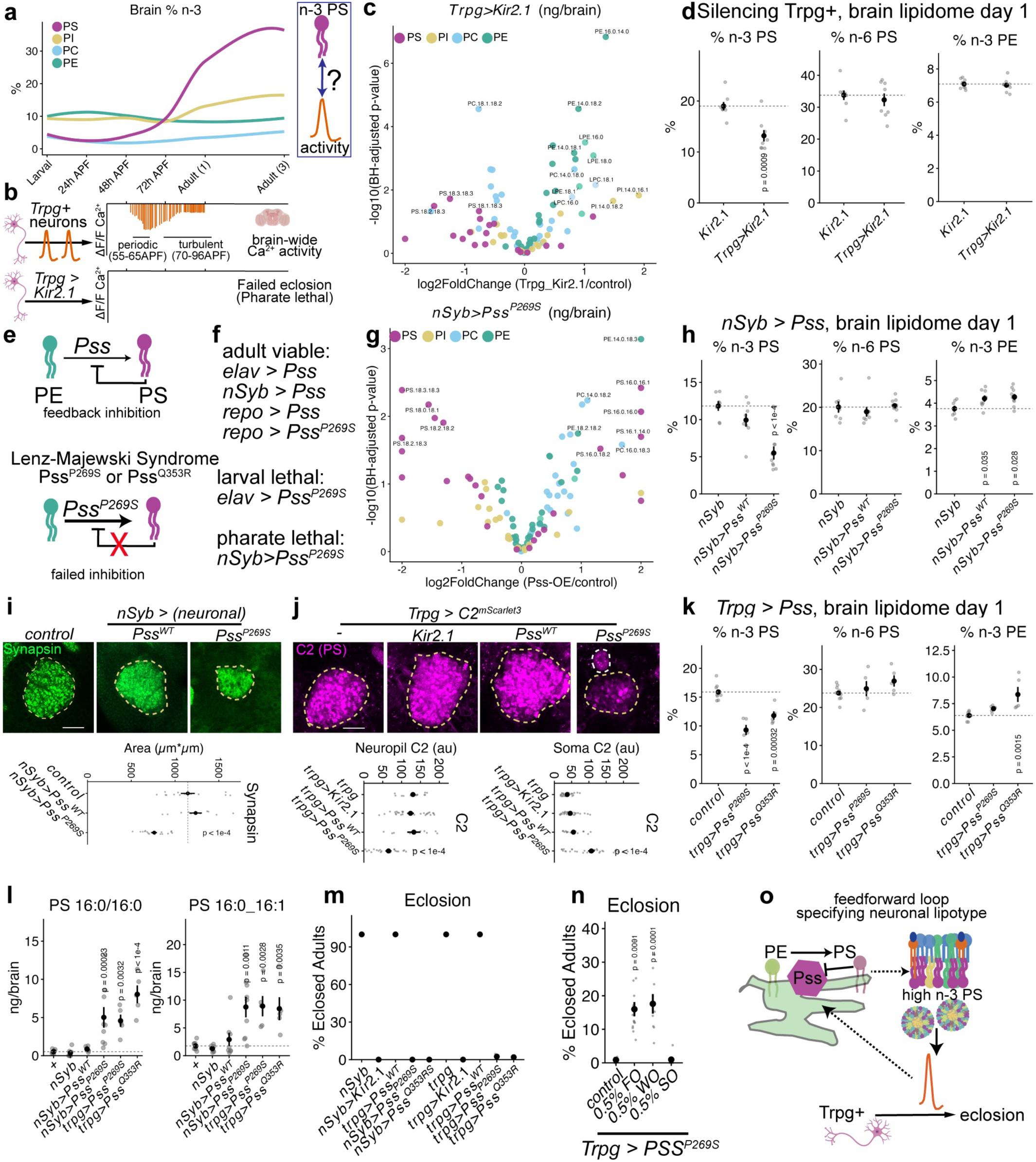
Neuronal activity and Pss feedback inhibition specify the functional neuronal lipotype. **a,** schematic of n-3 PS accumulation during development concordant with increased neuronal activity (orange lines). **b,** Trp*γ*+ neuronal activity templates brain-wide Ca^2+^ patterns from 55h APF; silencing Trp*γ* neurons abolishes Ca^2+^ and blocks adult eclosion. **c,** volcano plot of day 1 GPLs between controls and activity-silenced brains (*Trpγ* > *UAS-Kir2.1*). n=8 tubes per genotype. Annotated species are p<0.05 by BH-adjusted student’s t-test, and >0.5 log2 fold-change. **d,** % n-3 PS is selectively reduced upon activity silencing (*Trpγ* > *UAS-Kir2.1*). n=8 tubes per genotype. p-values indicate BH-corrected pairwise t-tests (two-sided); mean and SEM are shown. **e,** schematic of Pss regulation via PS-mediated feedback inhibition, and Lenz-Majewski Syndrome (LMS) variants, which block feedback inhibition. **f,** viability when inducing WT Pss or the analogous LMS variant Pss^P269S^ (P322S in *Drosophila*). **g,** volcano plot of day 1 GPLs between control and Pss^P269S^ brains (*nSyb > Pss^P^*^269^*^S^*). n=8 tubes per genotype. Annotated species are p<0.05 by BH-adjusted student’s t-test, and >0.5 log2 fold-change. **h,** % n-3 PS is selectively reduced in the brain upon *Pss^P^*^269^*^S^* induction. n=8 tubes per genotype. p-values indicate BH-corrected pairwise t-tests (two-sided); mean and SEM are shown. **i,** Synapsin staining in the mushroom body calyx of controls (n = 13 calyxes) vs *Pss^WT^* (n =11 calyxes) or *Pss^P^*^269^*^S^* (n=18 calyxes); scale bar = 20µm. **j,** C2 induced by *Trpγ-GAL4* in control (n=15 calyxes), *Kir2.1* (n=26 calyxes), *Pss^WT^*(n =9 calyxes) or *Pss^P^*^269^*^S^* (n=23 calyxes), with quantification of neuropil C2 (dashed yellow) versus soma C2 (dashed white). **k,** % n-3 PS is selectively reduced upon *Pss^P^*^269^*^S^*(n= 5 tubes) or *Pss^Q^*^353^*^R^*(n=5 tubes) induction with *Trpγ-GAL4* compared to controls (n = 8 tubes). p-values indicate BH-corrected pairwise t-tests (two-sided); mean and SEM are shown. **l,** PS 16:0/16:0 and PS 16:0_16:1 levels (ng/brain) are increased in Pss-LMS variants compared to controls. p-values indicate BH-corrected pairwise t-tests (two-sided); mean and SEM are shown. **m,** frequency of adult eclosion for the given genotypes. Dots represent mean of >3 independent crosses. **n,** rescue of failed eclosion in *Trpγ > Pss^P^*^269^*^S^* (n= 6 crosses) by supplementing diets with 0.5% flaxseed oil (FO; n = 14 crosses) or 0.5% walnut oil (WO; n = 9 crosses), but not with high-oleic (18:1) sunflower oil (SO; n = 6 crosses). **o,** model for n-3 PS specification and function in neurons via a feedforward loop combining neuronal activity and regulated Pss activity.

How might neuronal activity regulate n-3 PS specification? Activity-induced transcription of lipoprotein receptors^80,81^ motivated us to screen for altered n-3 PS levels in mutants for lipoprotein receptors (*LpR1;LpR2*), lipocalin homologs of APOD (*GLaz,* a glial lipocalin, and *NLaz*), and fatty acid binding proteins (*fabp* and *Obp44a*^82^). Of these mutants, *LpR1^Δ^LpR2^Δ^* and *GLaz^Δ^* most significantly, but pleiotropically, reduced PUFA GPLs, with distinct impacts on chain class: n-6 fatty acids were strongly reduced by *LpR1^Δ^LpR2^Δ^*, while n-3 fatty acids were reduced most by *Glaz^Δ^* (Extended Data Fig. 9g). While *fabp^Δ^* hypomorphs weakly impacted chain remodeling, *fabp^KD^* in developing glia and neurons severely disrupted glial and synaptic development, with strong and nonselective effects on all n-3 and n-6 chains that may be secondary consequences of failed glial morphogenesis (Extended Data Fig. 9e-f). Thus, while *LpR1/LpR2* and *GLaz* are important for essential fatty acid accretion in the adult brain, and *fabp* is required for glial morphogenesis, these pathways appear unlikely to confer specificity for n-3 PS production, similar to the lysosomal or dietary manipulations that globally affect all GPLs.

We next tested if membrane contact site and lipid transfer proteins might enable n-3 PS accretion, including PDZD8, an ER-lysosomal contact site protein critical for activity-dependent synaptic growth^83^, two bridge-like lipid transporters implicated in brain development (tweek^64^ and Atg2^84^), and Serinc, an activity-regulated ER incorporator of serine into PS^85^. None of these manipulations lowered n-3 PS, although *Atg2* perturbations reduced n-3 and n-6 PE and PC (Extended Data Fig. 9h-i). We also tested for altered ER-lysosomal contact in *Trpγ>Kir2.1* animals by Airyscan confocal imaging of the ER marker Bip-GFP and LysoTracker. While the overall volume of the ER and lysosome was similar between *Trpγ>Kir2.1* silenced animals and controls, we noted increased overlapping Lysosomal-ER signals in *Trpγ>Kir2.1*, with the ER locally adopting a denser, circular morphology that coincided with LysoTracker (Extended Data. Fig. 9j, arrows). Thus, silencing neuronal activity appears to alter the subcellular relationship between the ER and the lysosome.

We thus refocused on the ER enzyme Pss as a potentially instructive node in determining neuronal PS chain composition. Pathogenic variants in PTDSS1 alter PS chain composition in osteoclasts and cause intellectual disability in Lenz-Majewski Syndrome^16,86^. These gain-of-function mutations disable feedback inhibition, in which PTDSS1 is inhibited by PS^16,87^ (Fig. 7e). This prompted us to test whether feedback inhibition on Pss was critical for n-3 PS accretion during neurodevelopment. We expressed fly analogs of *Pss^WT^*, *Pss^P^*^269^*^S^*, or *Pss^Q^*^353^*^R^* in developing neurons, and found that while *elav-*driven *Pss^P^*^269^*^S^* or *Pss^Q^*^353^*^R^* was larval lethal, *nSyb-*driven adults survive to pharate stages, similar to *Trpγ>Kir2.1* animals (Fig. 7f). Interestingly, glial (*repo*) driven *Pss^P^*^269^*^S^* animals survived to adulthood, suggesting that neurons are less tolerant of failed Pss feedback inhibition. Remarkably, in *nSyb>Pss^P^*^269^*^S^*but not *nSyb > Pss^WT^* animals, n-3 PS was selectively reduced compared to n-6 PS and other n-3 GPLs (Fig. 7g-h, Extended Data Fig. 9k). Total PS levels were not increased (Extended Data Fig. 9c), similar to in Lenz-Majewski syndrome patients^16^. Moreover, *nSyb>Pss^P^*^269^*^S^*significantly reduced Synapsin area in the mushroom body calyx (Fig. 7i); this defect was not observed with *repo > Pss^P^*^269^*^S^* and was partially rescued by flaxseed oil feeding (Extended Data Fig. 9q). Thus, regulation of Pss activity via feedback inhibition is essential for both accumulating n-3 PS and proper neurodevelopment.

We next tested whether introducing pathogenic Pss variants into Trpγ+ neurons would alter PS distribution and Trpγ+ neuronal circuit function. First, we observed that the C2 sensor, though unchanged by *Pss^WT^* induction, was disrupted by *Trpγ>Pss^P^*^269^*^S^*, with significantly lower neuropil levels and higher cytosolic expression in cell soma, suggesting that aberrant PS chain composition could alter the subcellular localization of PS (Fig. 7j). Indeed, lipidomics of *Trpγ>Pss^P^*^269^*^S^* or *Trpγ> Pss^Q^*^353^*^R^* brains revealed a striking and selective reduction of n-3 PS compared to other n-3 and n-6 GPLs, as well as an accumulation of C16 saturated PS species, which also accumulated in *nSyb> Pss^P^*^269^*^S^*(Fig. 7l) but not in *Trpg>Kir2.1* silencing regimes (Extended Data Fig. 9m). Thus, Pss feedback inhibition from *within* the activity-templating Trpγ neuronal population is essential for n-3 PS accretion during development. Strikingly, *Trpγ> Pss^P^*^269^*^S^* and *Trpγ> Pss^Q^*^353^*^R^* animals also failed to eclose (Fig. 7m), suggesting that Pss feedback inhibition is crucial for Trpγ circuit function. Remarkably, this defect is sensitive to the presence of dietary n-3, as the pharate lethality of *Trpγ>Pss^PS^*^269^*^S^* was partially rescued by n-3 oils, whereas n-9 supplementation (high oleic sunflower oil) failed to rescue adult eclosion (Fig. 7n). While dietary n-3 supplementation also could rescue eclosion in *Trpγ>Pss^Q^*^353^*^R^*, it did not rescue eclosion when the *Trpγ* circuit was silenced by *Kir2.1*, despite n-3 chains accumulating in these brain (Extended Data Fig. 9n-o). We did not observe large LysoTracker accumulations in *Pss^P^*^269^*^S^* or other manipulations that altered n-3 chains in the brain, suggesting that these defects are not driven by lysosomal dysfunction (Extended Data Fig. 9p). Thus, not only does developmental neuronal activity govern n-3 PS accumulation, but proper PS chain composition within the Trpγ+ circuit appears indispensable for proper circuit function, n-3 PS accumulation, and adult eclosion. These data imply a feedforward loop between n-3 PS synthesis and neuronal activity (Fig. 7o), gated in part by PS feedback inhibition in the ER. In this model, the adult neuronal lipotype is furnished with the necessary substrates via dietary essential fatty acids and lysosomal catabolism of GPLs, before being specified by reinforcing Pss and neuronal activities (Fig. 8a).

**Fig. 8:**
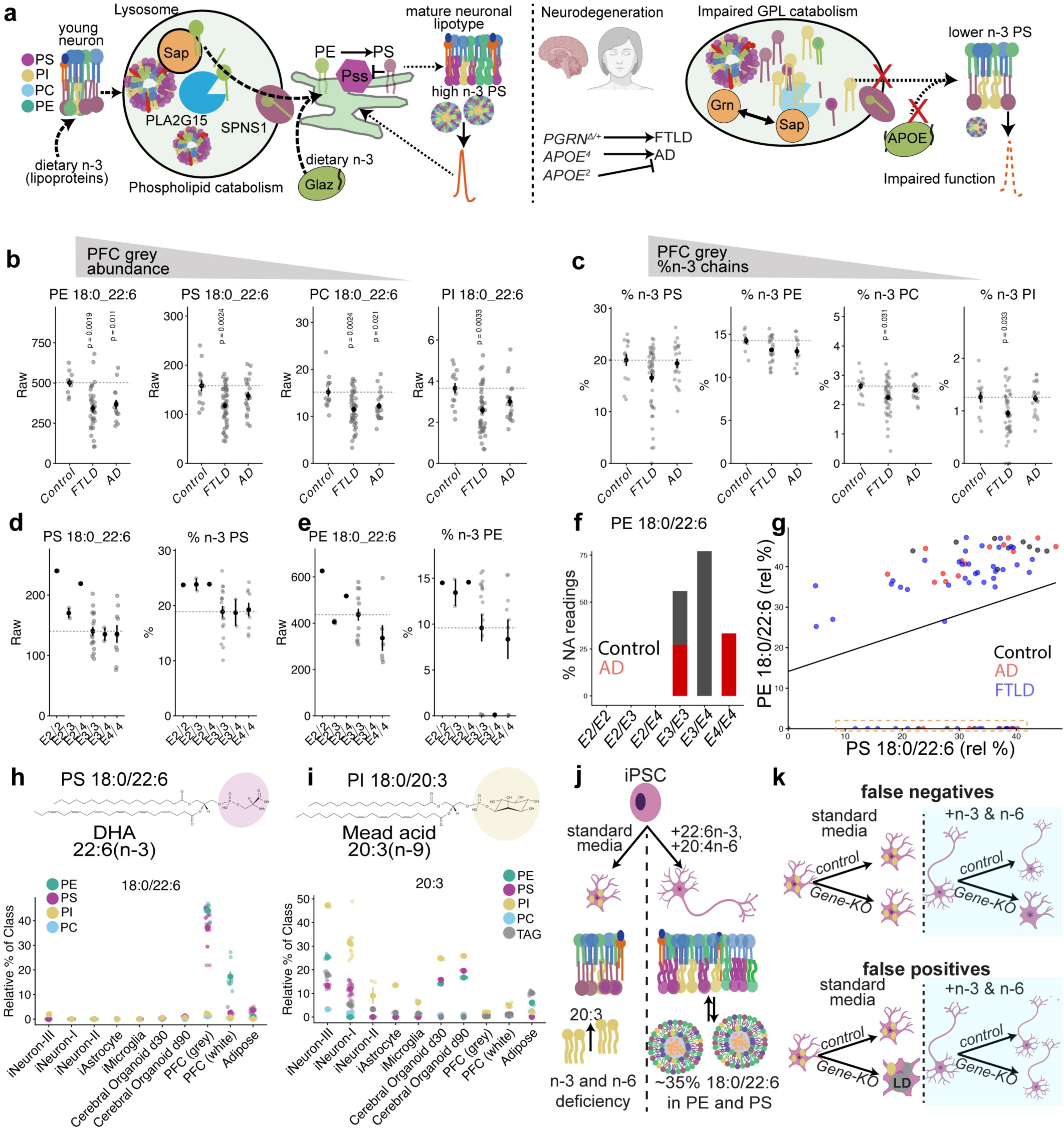
n-3 PS is disrupted in neurodegenerative disease but absent from *in vitro* models. **a,** model for how diet, neuronal activity, lysosomal catabolism, and PS synthesis generate mature neuronal lipotypes, and how neurodegenerative diseases (right, frontotemporal lobar dementia (FTLD) and Alzheimer’s (AD) diseases) may disrupt this process and impair n-3 PS accumulation and neuronal function. **b,** GPLs containing 18:0/22:6(n-3) raw data from Neurolipid Atlas^41^ prefrontal cortex (PFC) grey matter from controls, FTLD, and AD. n (patient samples) = 13 control, 46 FTLD, and 20 AD patients. **c,** % n-3 chain abundance from patients in b. p-values represent BH-corrected two-sided t-tests against control (dashed line). **d,** analysis of AD and control cohort stratified by APOE allele for PS 18:0/22:6 and % n-3 PS in grey PFC. Sample size (patients): E2/E2 =1; E2/E3=2; E2/E4=1; E3/E3 =18; E3/E4=2; and E4/E4 = 9. **e,** PE 18:0/22:6 and % n-3 PE in grey PFC from same patients as in d. **f,** proportion of 0/NA values for PE 18:0/22:6 in the Neurolipid atlas (red = AD, black = control), stratified based on APOE status. **g,** correlation plot for PE 18:0/22:6 and PS 18:0/22:6 in the Neurolipid Atlas (control = black, FTLD = blue, AD = red). Samples with NA values for PE are boxed in orange. **h**, relative % 18:0/22:6 across GPLs in human iPSC-derived cells from three independent labs^41,96^, cerebral organoids^99,100^ at two timepoints, human PFC grey or white matter^41^, and human adipose tissue^42^. **i**, relative % distribution of 20:3 chains (mead acid, n-9) across GPLs and triacylglycerols (TAG, grey). **j,** diagram of essential fatty acid deficiency triggered by *in vitro* conditions. **k,** essential fatty acid deficiency may yield false negatives or false positives. Data are represented by mean and SEM.

### n-3 PS is disrupted in neurodegenerative disease but is absent from *in vitro* models

Given n-3 fatty acid enrichment in human grey matter PS and PE (Fig. 1-2), we hypothesized that in neurodegeneration these highly abundant lipids may be altered. Indeed, frontotemporal dementia (FTLD) patients were recently reported to have decreased PUFA levels^88^. Intriguingly, lysosomal Progranulin (PRGN) mutations cause FTLD^89,90^, and PRGN regulates PSAP trafficking and function^91^. Moreover, a major risk factor for Alzheimer’s disease (AD), apolipoprotein E (APOE), carries PUFAs in lipoproteins^92,93^. Thus, mutations underlying FTLD and AD may alter lysosomal GPL catabolism, disrupting PUFA PS and driving neuronal dysfunction (Fig. 8a). To test this, we mined the recently released Neurolipid Atlas for lipidomics data from controls, FTLD, and AD patients^41^. Strikingly, PS 18:0/22:6 and PE 18:0/22:6, the two major n-3 brain GPLs, were decreased in FTLD patient grey matter, as were rarer PI and PC 18:0/22:6 lipids (Fig. 8b). While FTLD patients thus exhibited reductions in n-3 GPLs in the brain (Fig. 8c), n-6 GPL levels were unchanged, thereby lowering the n-3/n-6 ratio across GPLs in FTLD (Extended Data Fig. 10a-b). In contrast to FTLD patients, AD patients displayed a milder loss of 18:0/22:6 lipids, with the strongest effects in PE in both grey and white matter (Fig. 8b-c; Extended Data Fig. 10c). Thus, in FTLD grey matter, major n-3 PS lipids are reduced.

Intriguingly, all patients containing the neuroprotective APOE2 variant contained higher PS 18:0/22:6, n-3 PS, and PUFA PS in grey but not white matter (Fig. 8d; Extended Data Fig. 10d-f), with strongest effects on n-3 chains in PS and PE (Fig. 8e; Extended Data Fig. 10e). In a separate study of AD patients (ROSMAP)^94^, PUFAs in PFC grey matter were higher in APOE2/2 than APOE4/4 carriers for PS, but not for PE (Extended Data Fig. 10g). Surprisingly, in the Neurolipid Atlas, a high frequency of NA readings occurred for PE 18:0/22:6 and PE 16:0/22:6, both in control and AD patients harboring APOE3/3, APOE3/4, and APOE4/4 alleles (Fig. 8e-f; Extended Data Fig. 10h; n=14/29). Notably, patients harboring APOE2 alleles did not display PE-22:6 loss (Fig. 8f and Extended Data Fig. 10h; n=4). The loss of PE-22:6 also occurred in FTLD patients, and did not correlate with reduced PS 18:0/22:6 in these samples (Fig. 8g), suggesting a specific loss of PE-22:6 within the sample. Söderberg et al.^95^ observed large reductions in PE-22:6 in frontal grey matter in AD patients (Extended Data Fig. 10i) but did not note zero values, whereas ROSMAP did not contain missing PE-22:6 values. Together, these data sets suggests that APOE and AD could modulate levels of major n-3 PS and n-3 PE lipids, and that PE-22:6 appears particularly labile in non-APOE2 carriers within certain datasets.

Could a lysosomal Progranulin-PSAP axis account for the n-3 deficit in FTLD patients? A CRISPR screen identified PSAP as a core regulator of oxidative stress in iPSC-derived neurons (iNeurons)^96^. However, in analyzing the *PSAP* lipidomes produced in this study, we found that PS 18:0/22:6 and other 22:6n-3 species were absent from all genotypes (Fig. 8h; Extended Data Fig. 10j). Strikingly, iNeurons instead accumulated 20:3 chains across multiple lipid families, most notably in PI but also in PS, PE, and triacylglycerols (TAGs; Fig. 8i). As humans lack the desaturases required to produce n-3 and n-6 fatty acids, this signature is consistent with essential fatty acid deficiency, where mead acid (20:3n-9) is produced through other desaturases^97,98^. Absent 22:6n-3 and elevated 20:3n-9 occurred across separate iNeuron lines and organoids^99,100^ (Fig. 8h-i), indicative of a pervasive lack of n-3 fatty acids spanning many laboratories. Could iNeurons supplied missing essential fatty acids produce n-3 PS? By mining longitudinal proteomics from iNeurons^101^ for PTDSS1 and the lysosomal GPL catabolic module, we found that PTDSS1, PLA2G15, SPNS1, and PSAP are present in differentiating iNeurons (Extended Data Fig. 10k), with PLA2G15 and PTDSS1 expression increasing throughout differentiation. Intriguingly, the DHA-preferring acyl-CoA synthetase ACSL6 also increased throughout iNeuron differentiation, although the top-expressed ACSL, ACL3, prefers MUFA substrates^102^. These proteomics data support that iNeurons are competent to activate DHA-bearing GPLs and produce n-3 PS via PTDSS1 if supplied dietary n-3 fatty acids. Currently, with essential fatty acids missing from iNeuron media, a major lipid hallmark of the brain is not recapitulated by *in vitro* models.

## Discussion

We identify an evolutionarily ancient lipid hallmark of active brains, a high n-3/n-6 PS ratio, that is driven by the developmental accumulation of n-3 PS in synapses. This conserved lipid signature is underpinned by lysosomal lipid catabolism in developing neurons and glia, which alongside dietary n-3 lipids furnishes essential substrates for specifying high n-3 PS via cooperative neuronal activity and PS biosynthesis (Fig. 8a). Underscoring the intimate relationship between n-3 PS and activity, silencing developmental activity selectively lowered n-3 PS levels, while introducing pathogenic Pss variants into developmental circuits compromised circuit function and n-3 PS levels. This interdependent relationship between activity and n-3 PS links the emergence of physiology with the neuronal lipotype.

Why is n-3 PS enriched in neurons? Early^103,104^ and modern^35^ studies of purified synaptic vesicles found that DHA(n-3) was present in 30-40% of PS and PE, thereby contributing to ∼15% of total synaptic vesicle lipids. Tightly controlled acyl chains distributed across GPLs may enable the precise membrane gymnastics required for rapid endocytosis and synaptic vesicle curvature^12,13^. Consistent with this biophysical constraint, variation in PS desaturation and length was lowest in the brain compared to other tissues in mice^38^. Intriguingly, the PS flippase ATP8A1 localizes to synaptic vesicles and facilitates high frequency SV mobilization^105^. Pointing to a privileged relationship between PS and n-3 chains in the brain, neuronal cell lines exposed to DHA more avidly accumulate PS compared to other cells^106,107^, whereas depriving developing rats of DHA reduced total PS in the brain cortex but not in the liver or adrenal glands^108^. In mice, DHA levels correlated with neuron growth and Synapsin levels^109^, and disrupting maternal DHA compromised neuron morphogenesis in *Xenopus* tadpoles^110^. We now show that loss of lysosomal *Sap-r* reduces dietary n-3 incorporation into brain PS and other GPLs, and exacerbates Synapsin and behavioral deficits in *Pss^KD^* animals. Alongside regulating synaptic vesicles, n-3 PS may also be an important determinant of synaptic plasma membrane order^14^ and ion flux. PS 18:0/22:6n-3 represents the third most abundant synaptic junction lipid^111^, and n-3 PS is abundant in other electrically excitable tissues that lack synaptic vesicles, including muscle (Fig. 1), heart^112^, and retina^50^. Consistent with n-3 GPLs regulating membrane channel function and cellular excitability^9^, n-3 fatty acid supplementation during neurodevelopment potently rescued seizures caused by gain-of-function mutations in a voltage-gated sodium channel^113–115^. Thus, n-3 PS may sculpt the functions of both synaptic vesicles and membrane ion channels.

Our work here points to a crucial anabolic role for the lysosome in furnishing the essential building blocks for neuronal lipotype specification. In particular, we delineate a Saposin-PLA2G15-SPNS1 axis that controls PUFA accumulation in adult brain GPLs. Saposins are fascinatingly modular due to lysosomal cleavage into discrete proteins with different lipid affinities^24,31^. While the cleaved domains of *Drosophila* Sap-r are unmapped, the separability of sphingolipid and GPL defects suggests distinct functional modules (Extended Data Fig. 6). Thus, although sphingolipids are clearly dysregulated in *PSAP*-driven diseases^116,117^, our work also emphasizes that GPL defects may occur in *PSAP* and other lysosomal perturbations. Indeed, Saposin B can bind and transfer GPLs^33^, Saposin D solubilizes PS *in vitro*^32^, and PSAP promiscuously interacted with a panel of bifunctional lipids that included four GPLs^118^. Notably, rare Saposin D variants are associated with Parkinson’s disease in certain ethnic popoulations^119–124^, and cathepsins B and D, which may cleave PSAP, are candidate risk loci for Parkinson’s disease^125–128^. Given our finding that Sap-r regulates brain PUFAs, and prior work linking murine PSAP to cardiolipin chain saturation^30^, Saposin and the lysosome may broadly tune fatty acyl chain composition in the brain.

In addition to PSAP, PLA2G15 and SPNS1 homologs were critical determinants of lysosomal GPL catabolism and PUFA PS accumulation (Fig. 5). Rare biallelic mutations in SPNS1 cause multiorgan disorder with likely cognitive impairment and developmental delay in some patients^129^, and *spin* mutants reduce presynaptic neurotransmitter release^130^, trigger immediate lipofuscin accumulation^68^, and impair reproductive behavior^131^. Neurodevelopmental disorders have not been linked to PLA2G15 variants, but deleting the *C. elegans* homolog causes severe developmental defects^132^. While *PLA2G15^KO^* ameliorated *NPC1* or *TFEB* induced pathologies in mice through elevating anionic lysosomal lipids^71,133^, this genetic manipulation also causes splenomegaly, surfactant accumulation, and phospholipidosis^73^. Here, we demonstrate that deleting fly *PLA2G15* causes severe GPL accumulation, adult lethality, and a loss of synaptic density and circuit function, suggesting critical neuronal roles for PLA2G15. Consistently, PLA2G15 transcript levels were positively associated with cognitive function in AD patients^134^, and both differentiating fly and iPSC-derived neurons dynamically increased PLA2G15. Thus, depending on the context, PLA2G15-mediated lysosomal lipid processing may exert beneficial or detrimental effects. Importantly, in contrast to targeting *PLA2G15*, the loss of *Sap-r* did not cause lethality, meaning that *Sap-r* does not abolish *PLA2G15* activity. While the simplest interpretation of our data suggests a PSAP-PLA2G15-SPNS1 lysosomal GPL catabolic module (Fig. 8a), these proteins could plausibly operate in separate lysosomes or cell types. Interestingly, deleting *PGRN*, which disrupts PSAP, reduced PS in fibroblasts, which we speculate may arise via impaired lysosomal GPL catabolism^135^. Thus, lysosomal PSAP-PLA2G15-SPNS1 may broadly control both synaptic and systemic GPL composition.

Alongside the lysosome, we identified the regulated activity of the PS synthase Pss as crucial for n-3 PS levels and circuit function (Figs. 6-7). While loss-of-function variants in mammalian PS synthases have not been frequently implicated in neurodevelopmental or neurodegenerative disorders, there may be redundancy between PTDSS1 and PTDSS2^136^. Recently, copy number variants in *PTDSS1* were linked to autism^137^, and a heterozygous *PTDSS1* allele was associated with mild-to-moderate developmental delay^138^. In contrast, blocking feedback inhibition of PTDSS1 (i.e. *Pss^P^*^269^*^S^*) causes Lenz-Majewski syndrome^16^, a systemic disease featuring intellectual impairments and altered PS chain composition^86^. We found that inducing *Pss^P^*^269^*^S^* in developing neurons compromises the specification of the neuronal lipotype, with failed n-3 PS synthesis and the accumulation of saturated PS (Fig. 7). Moreover, *Pss^P^*^269^*^S^* perturbed developing circuit physiology in a manner sensitive to dietary n-3 chains. Reciprocally, silencing spontaneous developmental activity reduced n-3, but not n-6, PS. These surprising findings reveal complex and bidirectional relationships between neuronal activity and n-3 PS. It will be exciting to test how activity might alter organelle membrane contact sites over neurodevelopment, and we detected changes in ER-lysosomal overlap in silencing conditions that may alter Pss biosynthetic output. As PS synthases are calcium-dependent^87,139^, calcium signaling downstream of developmental activity could couple differentiation to PS biosynthesis, and changing membrane potential can alter PS clustering and intracellular signaling^140^. Notably, blocking developmental calcium alters synapse numbers in a cell-type specific manner^77^, *Sap-r* mutants have altered synaptic densities and calcium responses in the visual system^26,59^, and PS metabolism is crucial for critical period pruning of olfactory glomeruli^6^. Thus, interdependent relationships likely exist between neuronal activity and lipid metabolism to precisely coordinate neuronal lipotype specification with appropriate synaptic physiology.

Lipotype acquisition in the adult brain also depends on dynamic interactions between developing neurons and glia, with neuron-glia metabolic coupling a common theme in this complex tissue^7,59,81,82^. Based on the nonautonomous nature of the *Sap-r^Δ^* lysosomal patches (Fig. 4), with dual requirements for Sap-r in both developing neurons and glia, we propose that non-autonomous rescue occurs when glia consume neuronal membranes and thus enable complementation of Sap-r exchanged between cell types^59^; as *Sap-r* depletion in neurons or glia autonomously disrupted lysosome size, continuous and pervasive Sap-r exchange appears unlikely. How might developing neurons then receive glial-processed lipids? Our LC-MS/MS screening of candidate lipoproteins and lipoprotein receptors revealed that the glial-derived lipocalin GLaz (APOD) was important for accreting n-3 chains across all GPLs, whereas the lipoprotein receptors LPR1 and LPR2 was more critical for n-6 chain levels (Extended Data Fig. 9g). In mice, the transporter Msfd2a is essential for n-3 accretion by non-selective uptake of all lysophosphatidylcholines and is also required for barrier function^141,142^. Thus, while lipid transporters and lipoproteins are critical for accreting essential fatty acids in synapses, additional mechanisms, both upstream and downstream of lipid transport into the brain, likely tune the selectivity of the passaged dietary lipids and the ensuing high levels of synaptic n-3 PS.

We identified developmental diet as essential for specifying the adult neuronal lipotype. Strikingly, the chains loaded in the developmental diet perdured for 50 days (>60% lifespan), whereas swapping PUFA-free animals to regular diets as juvenile adults failed to accumulate equivalent levels of PUFA PS by day 50 (Fig. 2). Thus, neurodevelopment may represent a critical period for defining the lipidomic architecture of the adult brain, consistent with persistent developmental lipid signatures in the adult mouse brain^45^. Preterm infants may thus have reduced accretion of maternal DHA^143^, and treating them with DHA supplements improved Bayley mental and psychomotor developmental scores in a double-blind randomized trial^53^. Studies of brain DHA in rats point to lowest turnover rates of DHA in PS, with a 56 day half-life that is further extended by reduced n-3 dietary regimes^144^. In these studies, dietary depletion was conducted while the brain was still developing (P21), as depletion at P60 failed to impact DHA levels in the brain, whereas heart, testis, and liver all lost DHA^145^. In rhesus macaques, prenatal dietary depletion of n-3 and then juvenile supplementation did not fully restore visual acuity associated with n-3 deficiency, further pointing to the possibility of a critical period in neuronal lipotype specification^146^. While we have focused on n-3 PS, there is an interdependent relationship, or balance, between n-3 and n-6 fatty acids. For example, excess dietary n-3 fatty acids severely reduced n-6 chains in the brain (Fig. 3e). Functions for synaptic n-6 GPLs, which can yield bioactive signaling molecules, should also be explored in the context of heterogeneous neuronal membranes.

The absence of the major n-3 fatty acid (DHA) *in vitro*, and the corresponding increase in 20:3n-9, is likely of fundamental importance for studying neurodegeneration in iPSC-derived neurons, as is increasingly common during the investigation of metabolic diseases^147^. Deprivations of these essential fatty acids in infant humans^52–55,148^, rhesus macaques^149–151^, rats^50,51^, or flies^47–49^ cause aberrant neuronal physiology and behavior, defects that we suggest arise from impairing n-3 PS levels in synapses. Currently, if control cultured neurons suffer an essential fatty acid deficiency, false negatives may occur that mask important biology (Fig. 8k). Similarly, metabolic rewiring to high 20:3 fatty acyl chains (in PI, TAG, and PS) may yield false positives that selectively occur in n-3 deficient backgrounds (Fig. 8k). As TAGs are major components of lipid droplets, and PI-derived lipids are versatile signaling molecules that define organelle identities^152^, many biological processes may be altered in current culture models. Supplementing *in vitro* models with missing essential fatty acids is a promising strategy to rectify aberrant lipotypes and more accurately model neurobiology, and may increase culture viability, maturity, and activity^39,153^ (Fig. 8j). Similarly, boosting neuronal activity in culture may enhance n-3 incorporation into PS, as silencing developmental neuronal activity reduced n-3 PS (Fig. 7).

Aberrant metabolic rewiring is also of potential concern to studies focused on APOE and lipid droplets in iMicroglia^154^ or iAstrocytes^155,156^. There is growing evidence linking APOE to altered lipid^157^ and PUFA metabolism: APOE4 decreased the flux of oxidized PUFAs from neurons^158^, fomented lipofuscin deposition via PUFA-containing sterol esters^92^, and decreased n-3 PUFAs in human cerebrospinal fluid^159,160^. In contrast, hAPOE2 mice increased DHA uptake in the brain^161^ and were protected against ferroptosis^93^. Moreover, LPE-DHA is reduced in symptomatic AD patients^94^, and APOE2 positively correlated with higher PUFA PS, and negatively correlated with the sporadic lack of PE-DHA (Fig. 8d-f). Future studies should examine how APOE interfaces with n-3 fatty acids in brain lipid metabolism. DHA-bearing lipids also appear important during aging, as blocking DHA activation in neurons by mutating *ACSL6* impairs memory and motor function in mice^162,163^.

This work illuminates how the developing brain specifies a unique lipotype. Lysosomal catabolism and dietary inputs furnish the lipid building blocks that are then selectively sculpted by developmental circuit activity and regulated PS synthesis to endow synapses with abundant n-3 PS. Perturbing these developmental processes causes immediate behavioral defects in adult animals, and developmentally loaded n-3 chains perdure for the majority of adulthood in the fly brain. Moreover, n-3 GPL levels are significantly reduced in FTLD grey matter and altered by APOE variants. Thus, the loss of n-3 PS could be an underlying mechanism in neurodegenerative or neurodevelopmental diseases caused by genetic or environmental disruptions to the catabolism, synthesis, interconversion, or transport of this abundant synaptic lipid that is essential for proper circuit function.

## Online Methods

### *Drosophila* genetics and molecular biology

*Drosophila* were reared on cornmeal-molasses food (LabExpress, R) or PUFA-free food (LabExpress, CDD) at 25°C on a 12-hour light/dark cycle at 50-70% relative humidity. For oil supplementation, walnut oil (Gefen, “100% natural”), flaxseed oil (Flora, organic), or sunflower oil (La Tourangelle organic, high oleic) were mixed into melted R food at 0.1% or 0.5% by weight. For *RNAi*, 29°C and *UAS-Dcr-2* were used to enhance knockdown. Crosses were set in biological triplicate and typically repeated months later to confirm stable phenotypes. The PS probe was constructed by fusing mScarlet3 to the N-terminus of a minimal C2 domain of mouse lactadherin isoform1; the control probe had five hydrophobic residues mutated to A^4^. *UAS-Sap-r::V5* consisted of Sap-r (RA isoform) with a c-terminal linker (AAAAQQQ) followed by V5(x3). Alfa-tagged PLA2G15 constructs were constructed by inserting the Alfa tag (SRLEEELRRRLTE) downstream of the signal peptide followed by a 2x(GGGGS) linker. For Pss transgenes, P322S and Q406R correspond to human P269S and Q353R. Plasmids were synthesized by Twist Bioscience from pJFRC7-20XUAS (Addgene #26220) and injected by Bestgene into *attp14* or *attp2*. Unless noted otherwise, newly eclosed female flies (juvenile flies, day 1) were used for lipidomics and confocal assays; *PLA2G15^ΔDf^*, *Trpγ > Kir2.1* or *Pss^P^*^269^*^S^* animals failed to eclose and were dissected at 96 hAPF. For qRT-PCR, RNA was isolated from five brains of day 1 females using the Arcturus PicoPure kit, and *act*, *tub*, and *Sap-r* quantified by SYBR green. Detailed primers, genotypes, and plasmids (including *PLA2G15* and *Pss* variants) are in Table S1. For climbing, newly eclosed flies were genotyped, then tested the following day (day 2) in the negative geotaxis assay by firmly tapping cohorts of 5-7 flies to the bottom of a clear vial, before recording the time and distance flies took to cross a 5cm threshold within 60 seconds (averaged over 3 trials per condition). For activity, day 2 male flies were loaded into the *Drosophila* Activity Monitor (TriKinetics), and total activity was computed from day 3 to day 8 of adulthood split across light and dark phases.

### Brain live imaging and fixed immunohistochemistry

LysoTracker (Thermofisher) imaging was conducted similar to^58^, with live brains dissected and immediately stained in a 1:200 dilution of LysoTracker red or green (in 1X phosphate buffered saline (PBS)) for one minute before washing in 1XPBS and immediate confocal imaging. Zeiss LSM900 with Plan-APOCHROMAT 40X (1.3 NA; oil) was used to obtain z-stacks at 1.5µm intervals and 2x digital zoom, before import to FIJI, where particle analysis via a custom macro computed LysoTracker area per individual brain optic lobe blind to genotype. For Synapsin or brp/nc82 staining, brains were fixed with 4% paraformaldehyde (diluted in 1XPBS) for 25 minutes^58^. Post-fixation, brains were washed and permeabilized for 45 minutes in 0.3%PBS-TritonX in 1% NGS and stained (Synapsin 3C11^164^ at 1:10; brp/nc82^165^ at 1:10; and GLUL(BCN148.1.2H5) at 1:20, antibodies from DSHB) and matched secondaries (Thermofisher). Mushroom body calyx z-stacks were obtained at 1.5µm resolution and 2x digital zoom on a Zeiss LSM900 with Plan-APOCHROMAT 40X (1.3 NA). Synapsin area was calculated from the z-slice with maximal mushroom body calyx diameter in FIJI (brains mounted posterior-apposed to coverslip). Data were exported to R Studio and analyzed by student’s two-sided t-test Benjamini-Hochberg corrected for multiple comparisons, with an assumed normal distribution of Synapsin and LysoTracker areas. For BIP-GFP imaging, a LMS980 with 63x oil lens was used at 8x digital zoom to capture cell soma from *Trpg+* control neurons or those expressing *Kir2.1*, co-stained with LysoTracker, using Airyscan (0.29µm z-interval). Soma were masked in FIJI to exclude non-expressing cell LysoTracker signals, and analyzed blind to genotype with a macro that computed volume of the ER (3D Manager) and binary segmentation overlap of lysosomal and ER channels per z-slice.

### Lipidomics

Brains were live dissected in 1XPBS (10/tube) in quadruplicate, frozen in 90% methanol added to 10% PBS (v/v) prior to Folch extraction and LC-MS/MS on an Agilent 6430 triple quadrupole for fatty acyl chain composition with EquiSplash standards (Avanti 330731)^58^. Detailed extraction protocol and LC-MS/MS parameters are included in Supplemental Methods. Double bond position for fatty acid chains was confirmed by GC-MS at OmegaQuant (see Supplemental Methods). For flies, n-3 = 18:3, and n-6 = 18:2. For mice and humans, n-3 = (22:6, 20:5, or 18:3), and n-6 = (20:4, 22:4, and 18:2). Muscle data from^43^ did not contain chain-resolved data, and C36-40 with 6 double bonds was treated as a proxy for n-3 (DHA), while C36-40 with 4 double bonds for n-6 (AA). % PUFA (and % n-3) were analyzed on per-chain basis (e.g., 18:0_22:6 is split into separate chains before computing n-3 and total PUFA levels). Developmental mouse data included timepoints from^38,39^, while all human developmental data was from^11^. For volcano plots and dot plots, student’s two-sided t-test with Benjamini-Hochberg correction was performed (abundant lipid species analyzed were normally distributed). For multi-genotypic comparisons (e.g., rescues), Tukey’s ANOVA was conducted.

### Electron microscopy

Electron Microscopy was conducted at the CU Anschutz EM core facility using the JEOL JEM-120i, 120 kV TEM. Day 1 female brains were dissected under 0.1 M, 7.4 pH Sorenson’s phosphate buffer fixed in 2% paraformaldehyde/ 2% glutaraldehyde (v/v in 0.1 M Sorenson’s phosphate buffer). Detailed methods are available in Online Methods. Lysosomes were quantified in neuronal soma in FIJI.

## Data and Code Availability

*Drosophila* lipidomics data is deposited in Table S2. All other data and code are available from the lead author upon request.

---

**Extended Data Fig. 1:**
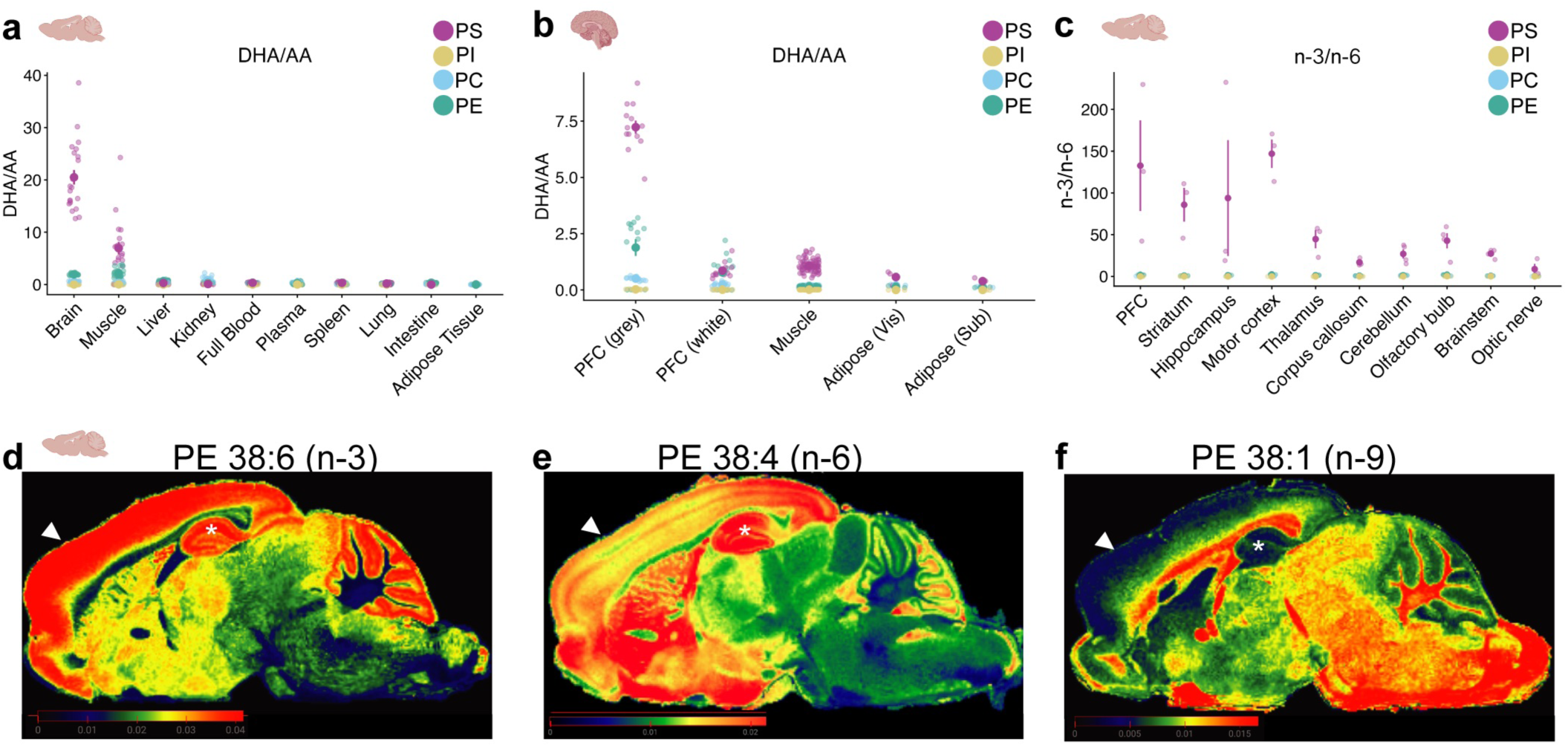
n-3 PS is abundant in neuronal regions. **a**, re-analysis of published data calculating n-3/n-6 ratio as DHA (22:6n-3) divided by AA (20:4n-6) across mouse tissues^23^ (from P60 mice, n = 23 tubes of brains; 23 muscle; 23 liver, 24 spleen, 23 kidney, 24 full blood, 24 lung, 24 adipose tissue, 24 intestine, and 24 plasma); **b**, n-3/n-6 ratio calculated as DHA (22:6n-3) divided by AA (20:4n-6) across human tissues^42,43,167^ (n = 13 PFC grey; 13 PFC white; 75 muscle samples; 3 visceral adipose tissue samples; and 3 subcutaneous fat samples). **c**, n-3/n-6 ratios of PS (purple) from mice^2^ are highest in synapse-dense areas such as prefrontal cortex (PFC). n = 3 for all regions except for n = 4 for brain stem, cerebellum, and olfactory bulb. **d,** re-analysis of MALDI-MSI^44^ of PE 38:6(n-3); **e,** PE 38:4(n-6); and **f,** PE 38:1(n-9) from a 6 month female mouse control brain (arrows, prefrontal cortex; *, hippocampus). Plotted data are represented by mean and SEM. Data are represented by mean and SEM. See Supplemental Table 1 for complete genotypes.

**Extended Data Fig. 2:**
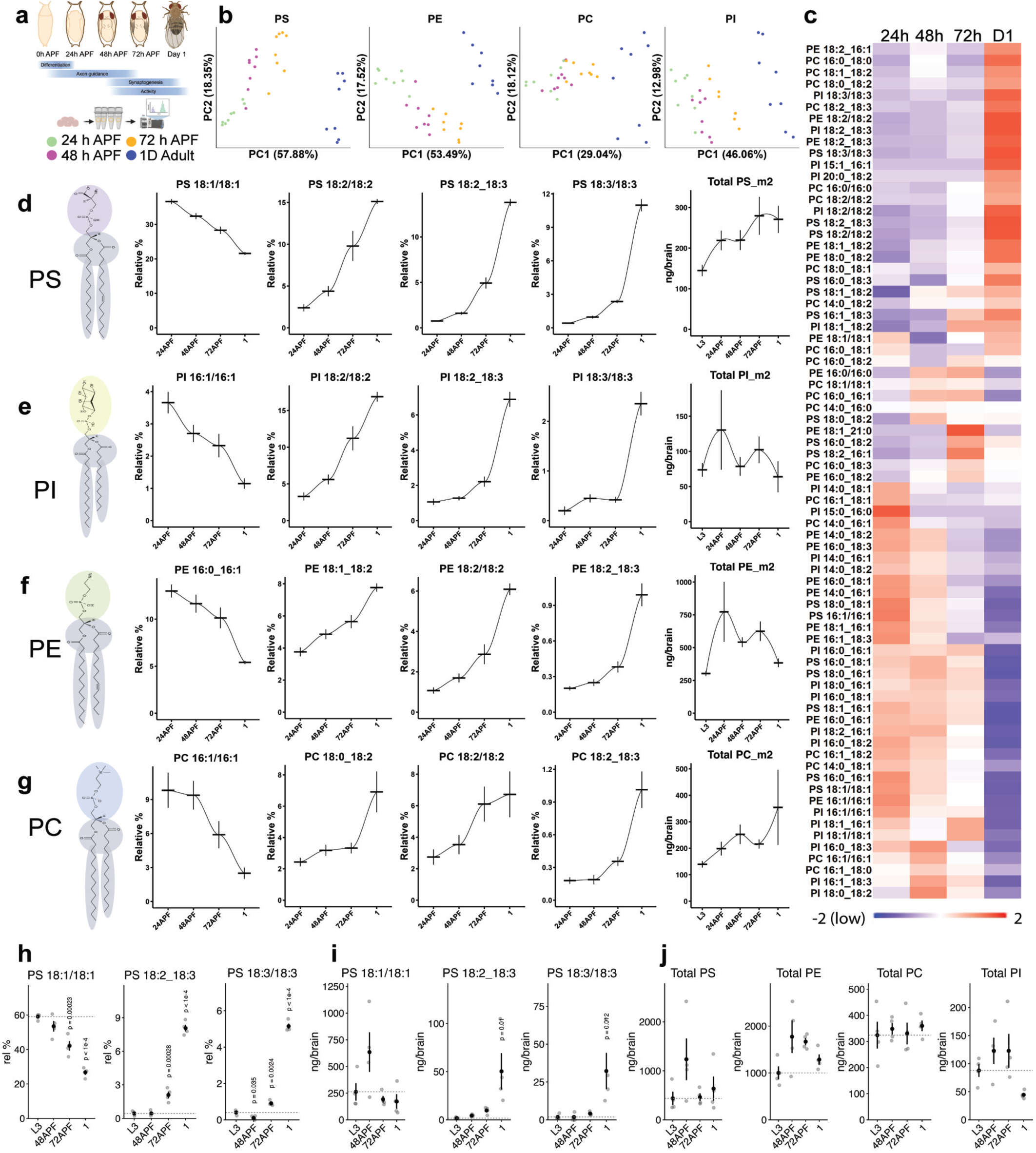
GPL remodeling during *Drosophila* brain development. **a,** schematic of *Drosophila* brain development. APF = hours after puparium formation. **b,** principal component analysis of major GPLs (phosphatidylserine (PS), phosphatidylinositol (PI), phosphatidylcholine (PC), and phosphatidylethanolamine (PE)) sampled every 24 hours. **c,** heatmap of z-scored abundances of major GPL. **d-g,** line plots of example species (relative % composition per class and total ng/brain distributions plotted over development), separated by lipid family. Dashes represent mean, error bars are SEM. **h-j,** independent lipidomics run over *Drosophila* neurodevelopment, with major PS species graphed (mean and SEM) as **h,** relative % composition or **i,** total ng/brain. **j,** total GPL ng/brain over neurodevelopment. n = 4 tubes per timepoint for h-j and 8 tubes per timepoint for a-g. See Supplemental Table 1 for complete genotypes.

**Extended Data Fig. 3:**
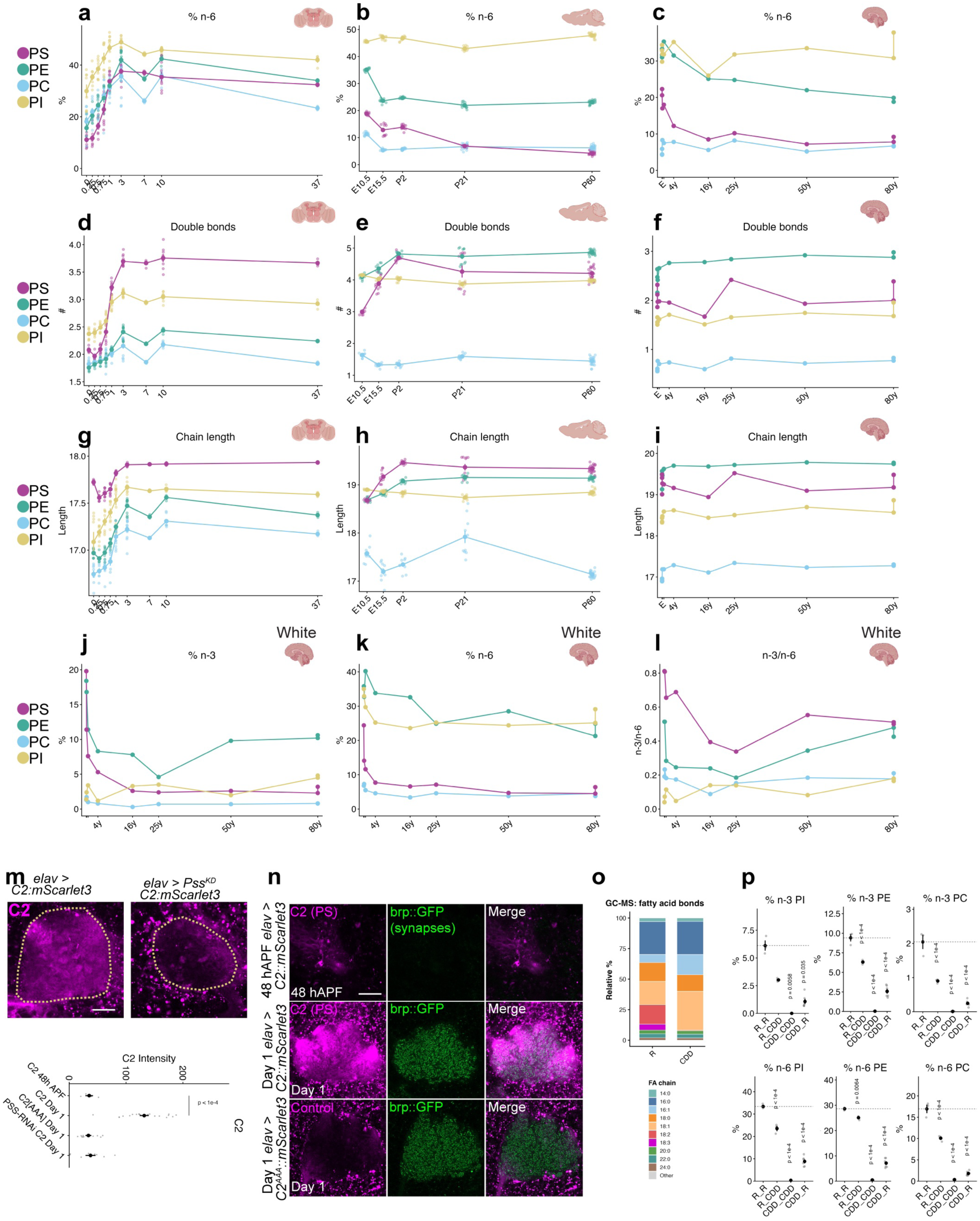
GPL remodeling across species, and dietary contributions in fly. **a,** % n-6 distribution of GPLs in brains dissected across development and adulthood in *Drosophila.* 0 = wandering larval stage L3, 0.25 = 24 hours after puparium formation (hAPF), 0.5 = 48 hAPF, 0.75 = 72 hAPF; 1-37 = day 1 to day 37 adults. n = 8 tubes for all ages except 7 and 37, where n = 4 tubes. **b,** GPL % n-6 distribution across mice brain development^38,39^; n = 10 tubes per all ages, except 23 tubes for P60. **c,** % n-6 GPL distribution in human cortex grey matter over age^11^ (n = 1 healthy control brain per listed age; E includes 12 weeks, 35 weeks, 38 weeks, 1 month, and 7 months; other ages indicate years). **d,** weighted fatty acyl double bond number across fly brain development and adulthood. **e,** weighted GPL double bond number across mouse brain development. **f,** weighted fatty acyl double bond number across human grey cortex brain development and adulthood. **g,** average fatty acyl chain length in GPLs across fly brain development and adulthood. **h,** average fatty acyl chain length in GPLs across mouse brain development. **i,** average fatty acyl chain length in GPLs across human grey cortex brain development and adulthood. **m,** neuronal expression of PS biosensor (*elav> C2^mScarlet^*^3^) imaged in control and neuronal *Pss^KD^*mushroom body calyx at day 1 (scale bar = 20µm). **n,** *C2^mScarlet^*^3^ imaged in mid-development (48h APF) and day 1 juvenile brains, imaged in mushroom body calyx with presynaptic active zones labeled by *brp::GFP* (green)*. C2^AAA^*control probe abrogates PS binding. Scale bar = 20µm. n > 10 calyxes per genotype. **o,** GC-MS analysis of precisely annotated double bond classes in brains from day 1 females of control or CDD diets (n= 4 tubes per condition). **p,** animals reared on control (R) or PUFA-free (CDD) diets were grown for 10 days into 1 day adults, then maintained on diets or swapped to reciprocal diets, before aging for 50 days and profiling brain lipid GPL distributions. % n-6 and % n-3 distributions in different GPLs are plotted. n = 3 tubes for R_R, 3 tubes for R_CDD, 3 tubes for CDD_CDD, and 6 tubes for CDD_R. p-values indicate BH-corrected student’s t-tests (two-sided) against control (R_R). Data are represented by mean and SEM. See Supplemental Table 1 for complete genotypes.

**Extended Data Fig. 4:**
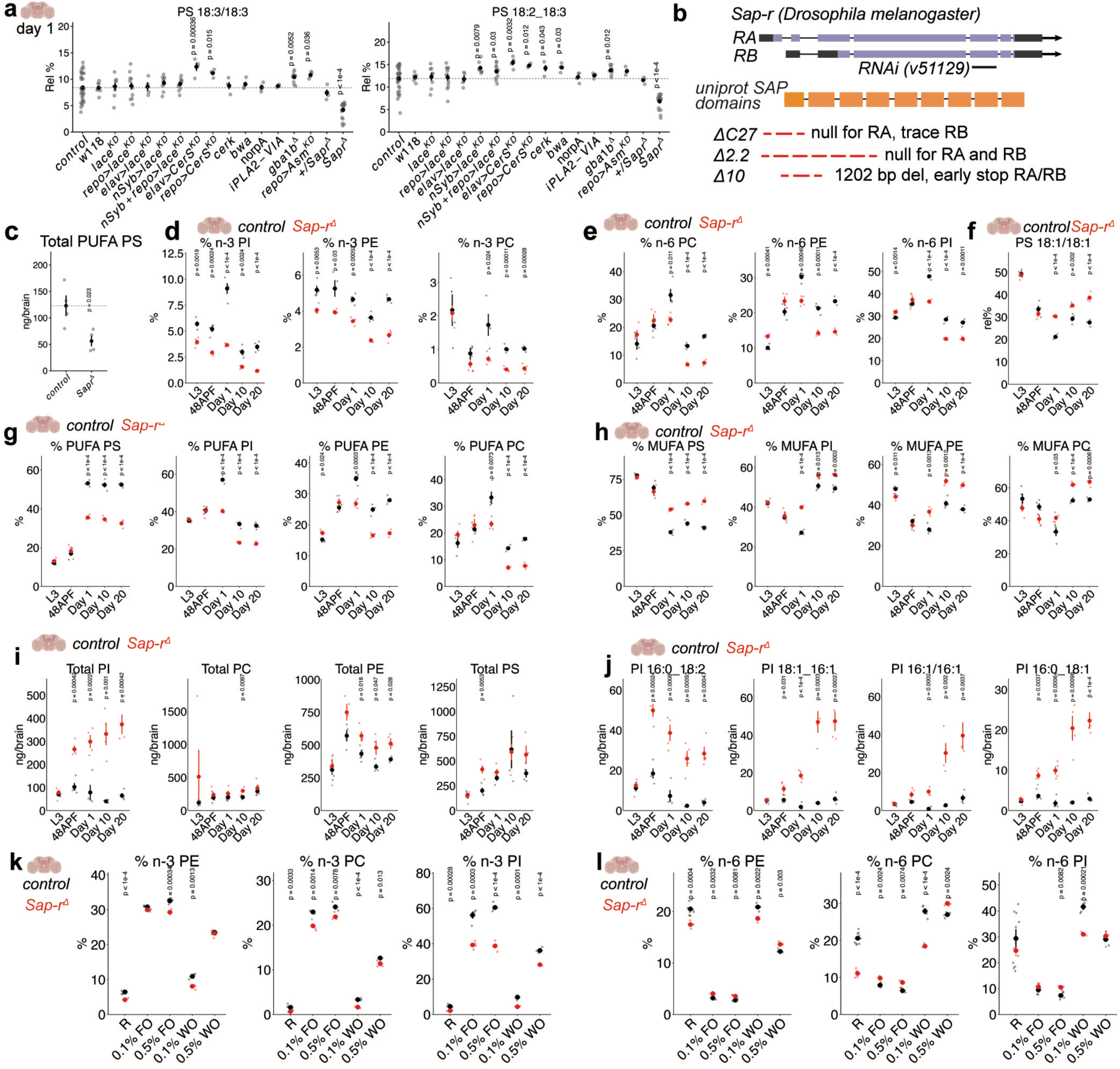
Saposin mediates PUFA accumulation in *Drosophila*. **a,** lipidomics of major brain PUFA PS species across control and mutant genotypes dissected at day 1 (n ≥ 4 tubes for lipidomics). p-values indicate BH-corrected student’s t-tests (two-sided). **b,** Sap-r locus and genetic reagents (dashed red line = deleted genomic DNA). **c,** total ng/brain levels of PUFA PS in control brains versus *Sap-r^Δ^* mutants at day 1. n = 4 tubes, p-values indicate BH-corrected student’s t-tests (two-sided). **d,** % n-3 chains in GPLs between controls and *Sap-r^Δ^* brains (red) across development (L3 = larval, 48APF = mid-development) out to 20 day adults; note that the x-axis (age) is a categorical variable. **e,** % n-3 chains in GPLs between controls and *Sap-r^Δ^* brains (red) across development. **f,** PS 18:1/18:1 relative % distribution in controls and *Sap-r^Δ^* brains (red) over development. **g,** % PUFA chains in GPLs between controls and *Sap-r* mutants (red) across development. **h,** % MUFA chains over development in controls and *Sap-r^Δ^* (red) brains **i,** total ng/brain for PI, PE, PS, and PC in controls and *Sap-r^Δ^* brains (red) over development. **j,** example PI species that accumulate in *Sap-r^Δ^*brains (red). **k,** control and *Sap-r^Δ^* (red) animals were reared on control (R), 0.1% flaxseed oil (FO), 0.5% FO, 0.1% walnut oil (WO), or 0.5% WO, and brains subjected to lipidomics at day 1, with % n-3 and **l,** % n-6 GPLs. For all lipidomics, n = 4 tubes/condition; p-values indicate two-sided t-test comparison of genotypes for a given diet or age. Data are represented by mean and SEM. See Supplemental Table 1 for complete genotypes.

**Extended Data Fig. 5:**
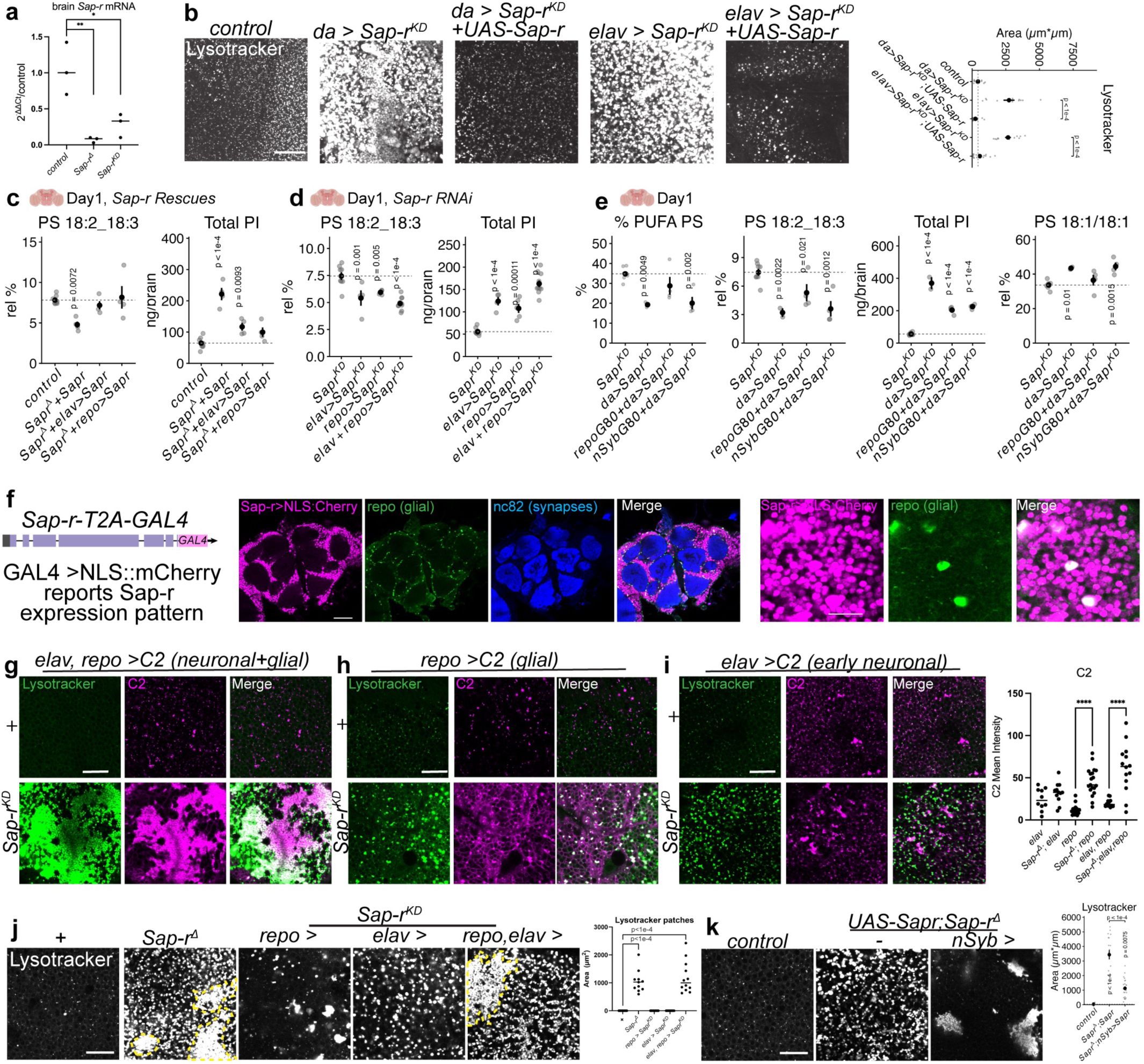
Cell-type specific analyses of Sap-r function. **a,** qRT-PCR analysis of control, *Sap-r^Δ^*, and *elav,repo>Sap-r^KD^* brains, normalized to control with internal normalization to tubulin and actin transcripts (n = 3 tubes of 5 brains per condition, p-value reports Tukey’s ANOVA (Prism). **b,** LysoTracker staining of day 1 optic lobes from *Sap-r^KD^* ubiquitously (*da-GAL4*) or in neurons (*elav-GAL4*) and rescue with synthetic *UAS-Sap-r* lacking exact sequence complementarity to the *RNAi* construct. Scale bar = 20µm, n > 10 optic lobes. **c-d,** day 1 brain total PI and PS 18:2_18:3 levels in cell-type specific rescue or knockdown experiments (*elav* or *repo* drivers). Lipidomics n: for rescue, 8 control tubes, 4 *UAS-Sap-r; Sap-r^Δ^* tubes; 4 *elav* > *UAS-Sap-r; Sap-r^Δ^* tubes; and 4 *repo UAS-Sap-r; Sap-r^Δ^* tubes; for knockdown, n=8 control tubes; 4 *elav* tubes; 5 *repo* tubes; and 10 *elav, repo* tubes. **e,** day 1 brain total PI and PUFA PS levels upon ubiquitous *Sap-r* knockdown (*da-GAL4*) with GAL4 suppression in maturing neurons (*nSyb-GAL80)* or glia (*repo-GAL80*). n = 8 tubes of control, 4 for *repo-GAL80+da-GAL4,* 4 for *nSyb-Gal80+da-GAL4,* and 2 for *da-*GAL4. **f,** endogenous transcriptional reporter for *Sap-r* (*Sap-r^T2A-GAL^*^4^) drives ubiquitous expression of *UAS-NLS::mCherry* (red), co-stained with anti-repo (green) to mark glia, account for only ∼10% of the fly brain. n=8 brains. **g-i,** day 1 optic lobe cortex stained for LysoTracker (Green) or PS (*C2^mScarlet^*^3^, magenta) in *Sap-r^KD^*driven in both neurons and glia (*elav,repo),* only glia (*repo)*, or only neurons (*elav*). Scale bar = 20µm, n > 10 optic lobes/genotype, p-value reports Tukey’s ANOVA (Prism). **j,** LysoTracker from optic lobes of cell-type specific *Sap-r^KD^* and controls, with LysoTracker patch area (yellow) quantified. n = 12 brains per genotype. p-value reports Tukey’s ANOVA (Prism) **k,** LysoTracker from optic lobes in control, *Sap-r^Δ^*, and *nSyb*+ *neuronal Sap-r* rescue. n (optic lobes) = control (8), *UAS-Sap-r; Sap-r^Δ^*(18); and *nSyb* > *UAS-Sap-r; Sap-r^Δ^* (23). p-values indicate BH-corrected student’s t-tests (two-sided). Data are represented by mean and SEM. See Supplemental Table 1 for complete genotypes.

**Extended Data Fig. 6:**
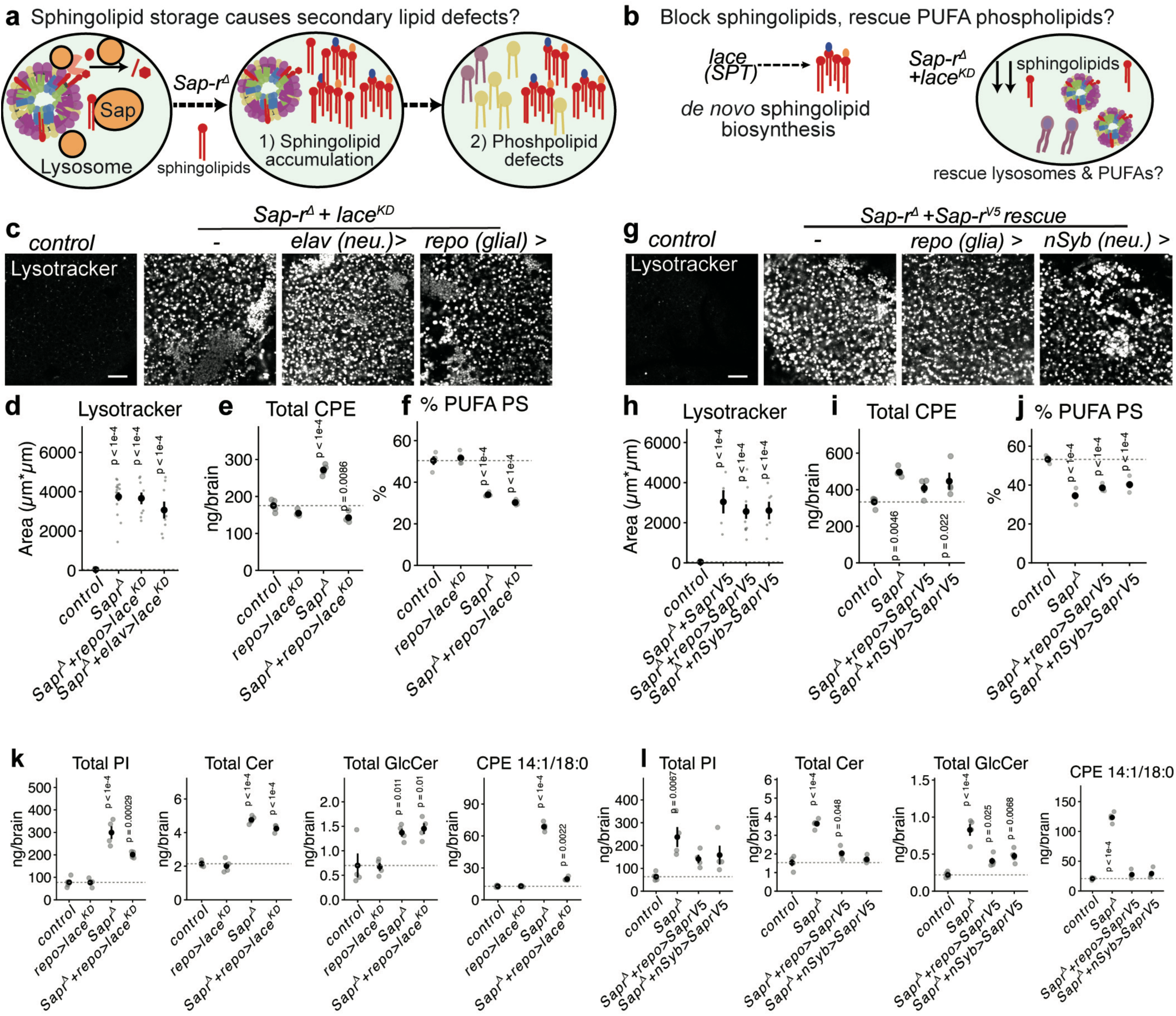
GPL deficits appear separable from sphingolipid accumulation in *Sap-r*. **a,** model for GPL defects arising secondarily to sphingolipid accumulation in *Sap-r^Δ^.* **b,** genetic method to block sphingolipid accumulation by disrupting *lace* (SPTLCB2). **c,** day 1 optic lobe soma stained for LysoTracker (white) in controls and *Sap-r^Δ^* mutants with *lace* knockdown in neurons or glia. Scale bar = 20µm. **d,** quantification of LysoTracker. n = 8 optic lobes for control; 16 *Sap-r^Δ^* ; 8 *Sap-r^Δ^* +*elav>lace^KD^;* and 9 optic lobes for *Sap-r^Δ^* +*repo>lace^KD^*; p-values represent BH-corrected two-sided student’s t-tests against control. **e,** quantification of the major sphingolipid ceramide phosphoethanolamine (CPE) in controls, *Sap-r^Δ^*, and glial *lace^KD^* in *Sap-r^Δ^*. **e,** PUFA PS quantification from genotypes in d. For lipidomics, n = 4 tubes for all genotypes; p-values represent BH-corrected two-sided student’s t-tests against control. **g,** day 1 optic lobe soma stained for LysoTracker (white) in controls and *Sap-r^Δ^*mutants expressing *Sap-r^V^*^5^ in neurons or glia. **h,** quantification of LysoTracker. n = 8 optic lobes for control; 5 *Sap-r^Δ^* ; 11 *Sap-r^Δ^* +*repo>Sap-r^V^*^5^; and 6 optic lobes for *Sap-r^Δ^* +*nSyb>Sap-r^V^*^5^. **i,** CPE in controls and *Sap-r^V^*^5^ rescues. **j,** PUFA PS in controls and *Sap-r^V^*^5^ rescues. **k-l,** total PI, Cer, GlcCer, and CPE 14:1/18:0 in controls, *Sap-*r*^Δ^*, and attempted rescue with glial *lace^KD^* or *Sap-r^V^*^5^ expressed in neurons (*nSyb-GAL4*) or glia (*repo-GAL4*). For lipidomics, n = 4 tubes for all genotypes; p-values represent BH-corrected two-sided t-tests against control. Scale bar= 20µm. Data are represented by mean and SEM. See Supplemental Table 1 for complete genotypes.

**Extended Data Fig. 7:**
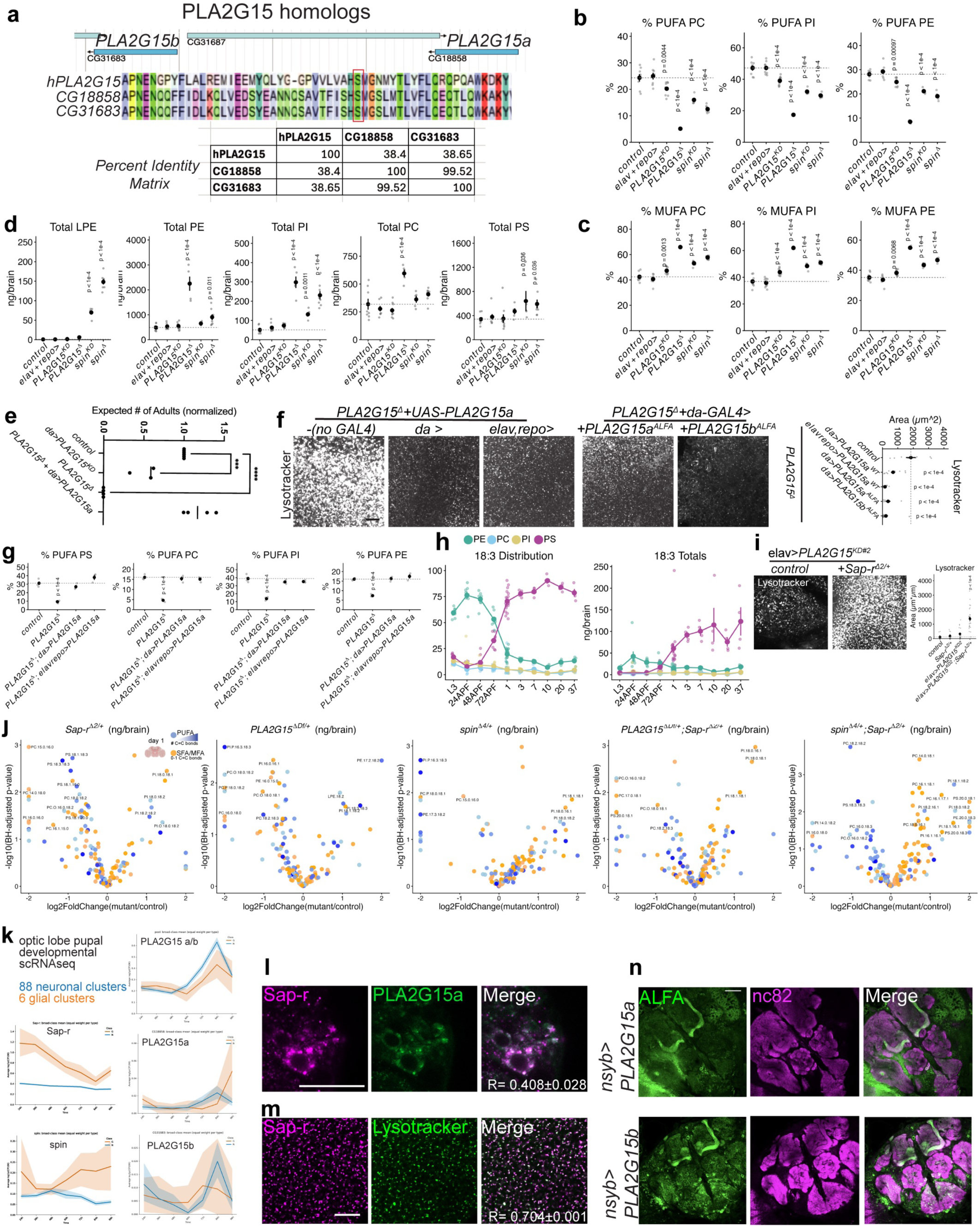
PLA2G15 and SPNS1 homologs mediate GPL remodeling. **a**, PLA2G15 homologs in fly (*CG18858 = PLA2G15a,* and *CG31683= PLA2G15b*) are neighboring duplications; the active site Serine is boxed in red and aligned with human PLA2G15 (hPLA2G15). **b,** % PUFA levels in day 1 brain from controls, *PLA2G15*, and *spin* manipulations. n = 8 wild-type tubes (*cs/iso*), 8 GAL4 control tubes (*elav-GAL4,repo-GAL4*/+); 9 *PLA2G15^KD^*tubes; 4 *spin^KD^* tubes; 5 *PLA2G15^Δ^* tubes; and 5 *spin^Δ^* tubes. p-value reports BH-corrected two-sided student t-tests. **c,** % MUFA per GPL class (same genotypes as b). **d,** total GPL and lysoPL levels in *spin*, *PLA2G15,* or control day 1 brains. p-values represent BH-corrected two-sided student’s t-tests against control**. e,** expected adult viability of >3 crosses for control (normalized to 1), ubiquitous *PLA2G15^KD^*, *PLA2G15^Δ^*, or *PLA2G15^Δ^*+ubiquitous rescue of *PLA2G15a.* **f,** LysoTracker staining of day 1 optic lobes from *PLA2G15^Δ^* with ubiquitously (*da-GAL4*) or neuronal+glial (*elav, repo-GAL4*) rescues of PLA2G15a, or with ubiquitous rescue of *PLA2G15^Δ^* via *PLA2G15a^ALFA^* or *PLA2G15b^ALFA^.* Scale bar = 20µm, n > 8 optic lobes, p-value reports BH-corrected two-sided student t-tests. **g**, brain lipidomics of *PLA2G15^Δ^* vs ubiquitous (*da*) or brain-restricted (*elav,repo*) *PLA2G15a* rescue. n = 3-4 tubes/genotype. **h,** lineplots of 18:3 chain % or ng distribution across PE (green), PS (magenta), PC (blue), and PI (yellow) in fly brains over development and adulthood. L3 = wandering larval stage L3, 24APF = 24 hours after puparium formation, and 1 to 37 = day 1 to day 37 adults. n = 8 tubes for all ages except 7 and 37, where n = 4 tubes. **i,** optic lobe LysoTracker in neuronal knockdown using a weaker *PLA2G15-RNAi* (KD2) in controls or heterozygous *Sap-r^Δ/+^.* Scale bar = 20µm. n= 16 control; 17 *Sap ^Δ/+^*; 19 *PLA2G15^KD^*^2^ ; and 26 *Sapr^Δ/+^ +PLA2G15^KD^*^2^ optic lobes, p-values report BH-corrected two-sided student t-tests. **j,** volcano plots from brain lipidomes of day 1 females for indicated control and transheterozygous genotypes (n = 5 tubes/mutant genotype, and 8 control tubes). **k,** reanalysis of scRNAseq optic lobe developmental dataset^61^ for glial (orange) and neuronal (blue) cluster expression of *Sap-r*, *spin*, pooled reads mapping to *PLA2G15a* and *PLA2G15b,* and unique reads mapping to *PLA2G15a* and *PLA2G15b*. **l,** fixed colocalization of *Sap-r^mScarlet^*^3^ (magenta) and *PLA2G15a^ALFA^* (green) driven neuronally by *nSyb-GAL4*. Scale bar = 10µm, n = 5 brains. **m,** live colocalization of *Sap-r^mScarlet^*^3^ (magenta; note that live retains more signal than fixed for lysosomal proteins in the fly brain) and LysoTracker (green). Scale bar = 20µm, n =5 brains. **n,** *nSyb-*driven expression of *PLA2G15a^ALFA^*, or *PLA2G15b^ALFA^*, stained for ALFA (green) or synapses (magenta). Scale bar = 50µm, n > 5 brains/genotype. Data are represented by mean and SEM. See Supplemental Table 1 for complete genotypes.

**Extended Data Fig. 8:**
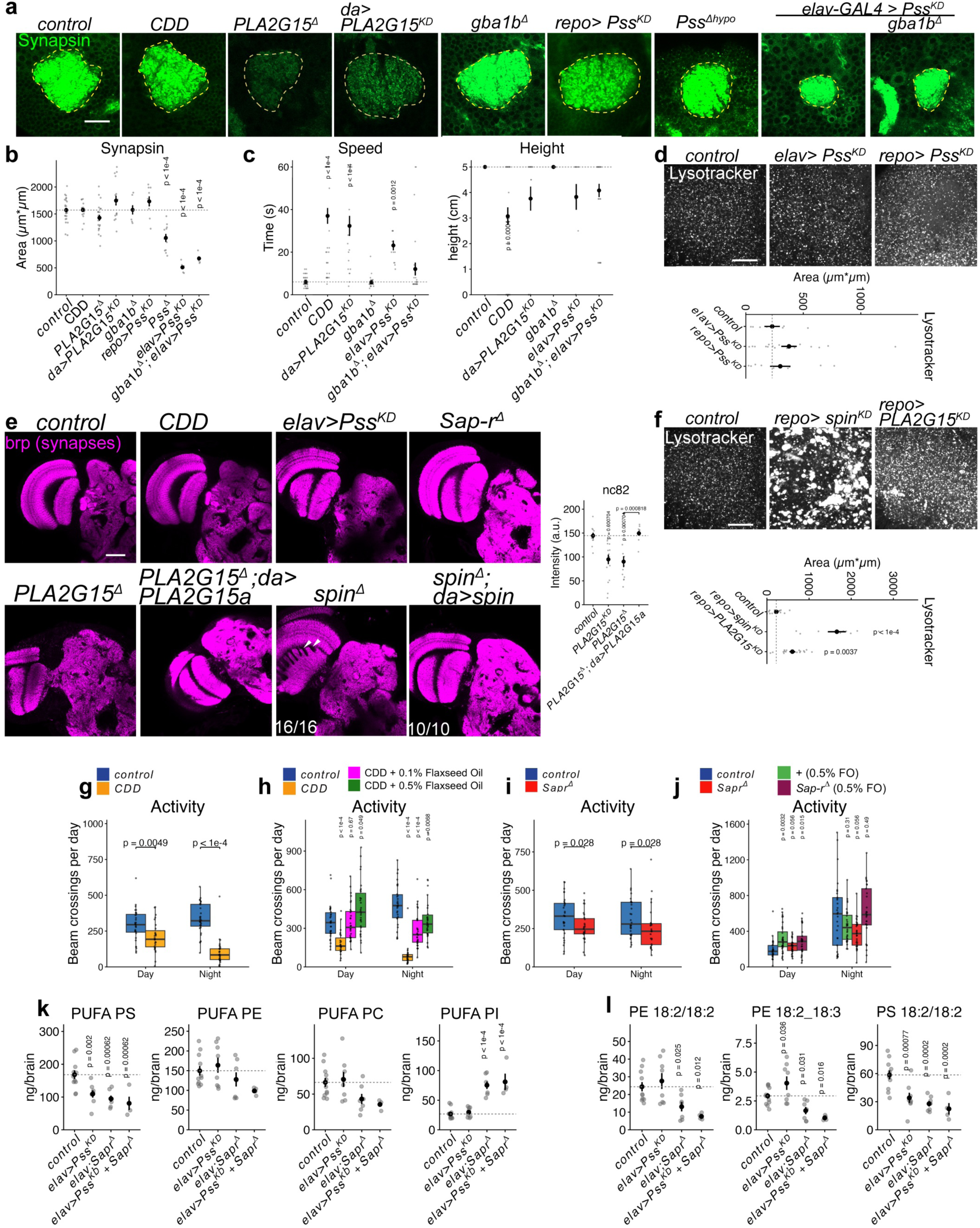
Neuronal PS biosynthesis and lysosomal mutant characterizations. **A-b**, day 1 mushroom body calyx (yellow outline) stained with Synapsin (green) in control (n=26 brains), CDD diet (n=16 calyxes), *PLA2G15^Δ^* (n=22 calyxes), *da> PLA2G15^KD^* (n = 16 calyxes), *gba1b^Δ^* (n = 7 calyxes), *Pss^hypomorph^* (n=11 calyxes), glial *repo>Pss^KD^* (n = 9 calyxes), neuronal *elav>Pss^KD^*(n=6 calyxes), and *gba1b^Δ^ + elav>Pss^KD^* brains (n = 6 calyxes); scale bar = 20µm. p-values represent BH-corrected two-sided t-tests against control. **c**, time and total distance in 5cm climbing assay (capped 60s) in day 2 females of: n = 44 flies (control, *isoD1*); 41 control flies reared on CDD; 28 *gba1b^Δ^* flies; 21 *PLA2G15^KD^* flies; 13 *elav >Pss^KD^* flies; and 28 *elav>Pss^KD^*+*gba1b^Δ^*flies. p-values represent BH-corrected two-sided t-tests against control. **d,** LysoTracker staining of day 1 optic lobes from neuronal (*elav*; n=19 optic lobes) or glial (*repo*; n =9 optic lobes) knockdown of *Pss*. Scale bar = 20µm, p-value reports BH-corrected two-sided student t-tests. **e,** day 1 brain synapses visualized by brp/nc82 (magenta) for indicated genotypes: lobula plate morphological abnormalities in *spin* mutants are indicated by white arrows. nc82 is quantified in mushroom body calyxes from control (n=10), *PLA2G15^Δ^*(n=12), *PLA2G15a* rescue (n=9), and *da> PLA2G15^KD^* (n=16). **f,** LysoTracker staining of day 1 optic lobes from glial knockdown of *spin* (n= 9 optic lobes) or *PLA2G15* (n=17 optic lobes) versus control (n = 16 optic lobes). Scale bar = 20µm, p-value reports BH-corrected two-sided student t-tests.**g,** day and nighttime activity (counts/fly) from male control (*isoD1)* flies developed from either regular food or CDD (PUFA-free), and then shifted at day 2 of adulthood to control food in activity monitors. Activity was quantified over the subsequent 5 days of adulthood (n=32 flies per genotype). p-value represents Mann-Whitney U test. **h,** same paradigm as f with 0.1% flaxseed oil (FO) or 0.5% FO supplementation of developmental CDD diets. n = 32 flies per genotype; p-value represents Mann-Whitney U test. **i**, activity profiles of control (*cs/isoD1*) and *Sap-r^Δ^*^10^*^/^*^22^ males (n = 32 flies per genotype). p-value represents Mann-Whitney U test. **j,** same genotypes as h, with 0.5% FO supplementation to developmental R diet. n = 32 flies per genotype; p-value represents Mann-Whitney U test. **k,** total PUFA GPLs (ng/brain) in control (11 tubes) versus *elav>Pss^KD^*(8 tubes) versus *elav;Sap-r^Δ^* (7 tubes) versus *elav>Pss^KD^*+ *Sap-r^Δ^*(4 tubes). **l,** same genotypes as d, plotting examples of individual PUFA PE and PUFA PS lipid species decreased by *elav>Pss^KD^*+ *Sap-r^Δ^*. p-values represent BH-corrected two-sided t-tests against control. Data are represented by mean and SEM. See Supplemental Table 1 for complete genotypes.

**Extended Data Fig. 9:**
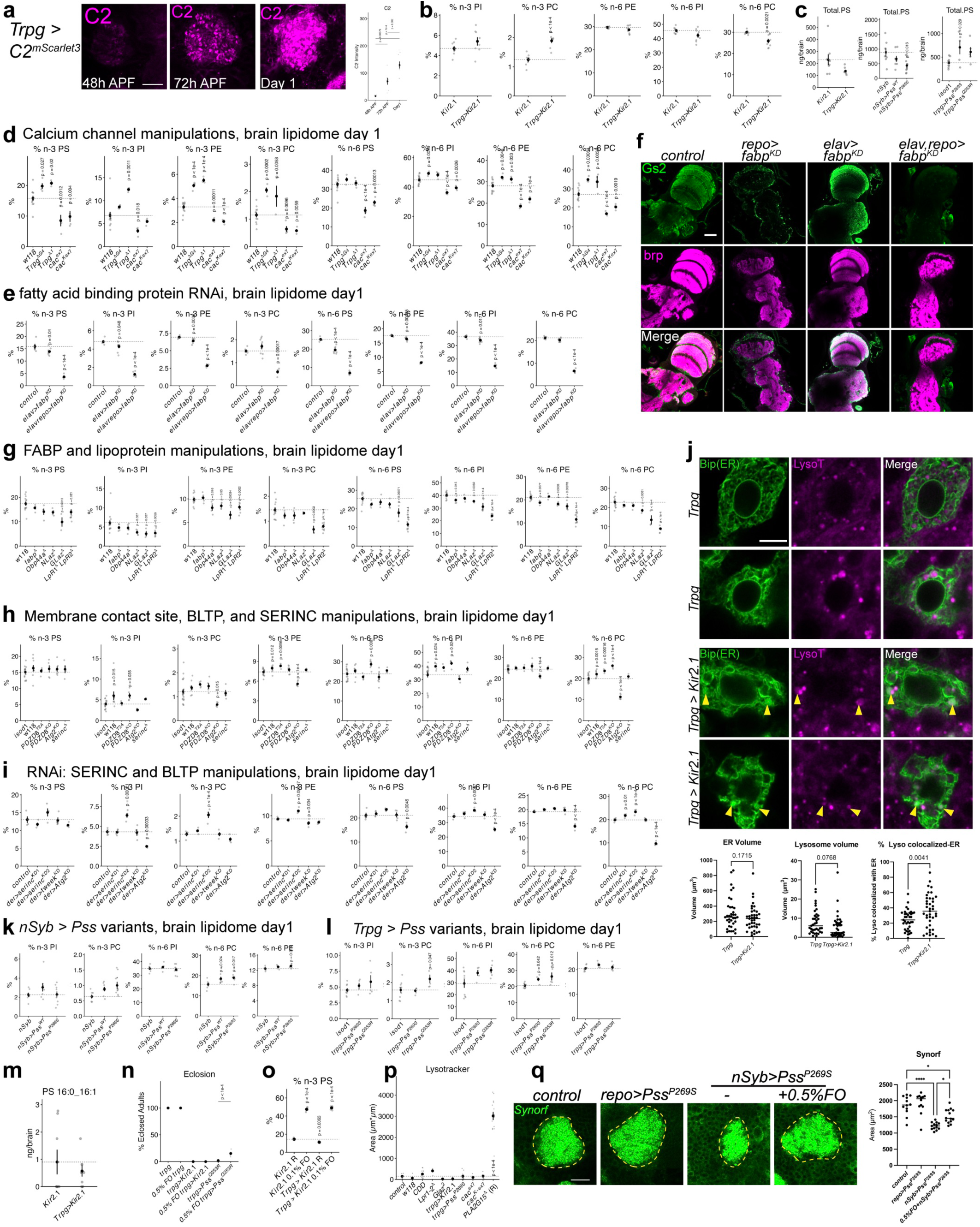
Mechanisms of n-3 PS specification in neurodevelopment. **a**, mushroom body calyx C2 driven by *Trpγ*-*GAL4* at 48hAPF, 72hAPF, and day 1. n > 7 calyxes per genotype. p-values represent BH-corrected two-sided t-tests against control. Scale bar = 20µm**. b**, % n-3 and n-6 GPL levels between controls and developmental neuronal activity silencing (*Trpγ>Kir2.1*). n=8 tubes per genotype. For this and all lipidomics data, p-values indicate BH-corrected pairwise t-tests (two-sided).**c**, Total PS/brain (day 1) in silencing and Pss gain-of-function manipulations (n = 8 tubes for control, *nSyb* and *Kir2.1* genotypes, and 5 tubes for *Trpγ>Pss* variants). **d**, %n-3 and %n-6 composition in GPLs from day 1 brains in controls vs *Trpγ* or *cac* mutants (n ≥4 tubes/genotype). **e**, %n-3 and %n-6 composition in GPLs from day 1 brains in controls vs *fabp RNAi* (n ≥4 tubes/genotype). **f**, staining for glial (Gs2+, green) and synaptic (brp/nc82, magenta) markers in control and *fabp-RNAi* brains. Scale bar = 50µm. Glial removal of *fabp* causes severe ultrastructural defects (loss of Gs2+ neuropil glia and malformed nc82; laser power was increased for nc82 signal in malformed brains to display defective architecture). n = 5 brains/genotype. **g**, %n-3 and %n-6 composition in GPLs from day 1 brains in controls vs lipoprotein mutants (n ≥3 tubes/genotype). **h**, %n-3 and %n-6 composition in GPLs from day 1 brains in controls vs bridge-like lipid transporter (BLTP), membrane contact site, and *serinc* mutants (n ≥4 tubes/genotype). **i**, %n-3 and %n-6 composition of GPLs from day 1 brains in controls vs BLTP and *serinc* knockdowns (n ≥4 tubes/genotype). **j,** *Trpγ* neurons expressing the ER marker Bip-GFP (green) co-stained with LysoTracker (magenta) in controls and silenced conditions (+*Kir2*.1) at 96 hAPF. Scale bar =5µm. n= 37 control cells and 40 *Kir2.1* silenced cells from >8 brains each. p-value indicates Prism unpaired t-test (two-sided). **k,** %n-3 and %n-6 composition of GPLs from day 1 brains in controls vs *nSyb-*driven *Pss* overexpression constructs (n =8 tubes/genotype). **l,** %n-3 and %n-6 composition in GPLs from day 1 brains in controls vs *Trpγ*-driven *Pss* overexpression variants (n≥5 tubes/genotype). **m,** total PS 16:0_16:1 (ng/brain) in *Trpγ*-silenced (+Kir2.1) brains. **n,** Adult eclosion frequency of listed genotypes and diets; eclosion failure of *Trpγ>Pss^Q^*^353^*^R^*, but not *Trpγ>Kir2.1*, can be partially rescued by feeding 0.5% Flaxseed Oil (FO). **o,** lipidomics of *Trpγ>Kir2.1* brains fed 0.5% Flaxseed oil reveals that dietary boosting of n-3 incorporation into PS can occur in neurodevelopmental silencing regimes. Data are represented by mean and SEM. **p,** Quantification of LysoTracker staining from day 1 optic lobes from control, calcium channel, and lipoprotein manipulations; *PLA2G15* is included as positive control (n ≥6 optic lobes/genotype). p-values represent BH-corrected two-sided t-tests against control. **q,** mushroom body calyx Synorf area for indicated genotypes, n > 11 calyxes per genotype. p-values represent Prism Tukey’s ANOVA. Scale bar = 20µm. See Supplemental Table 1 for complete genotypes.

**Extended Data Fig. 10:**
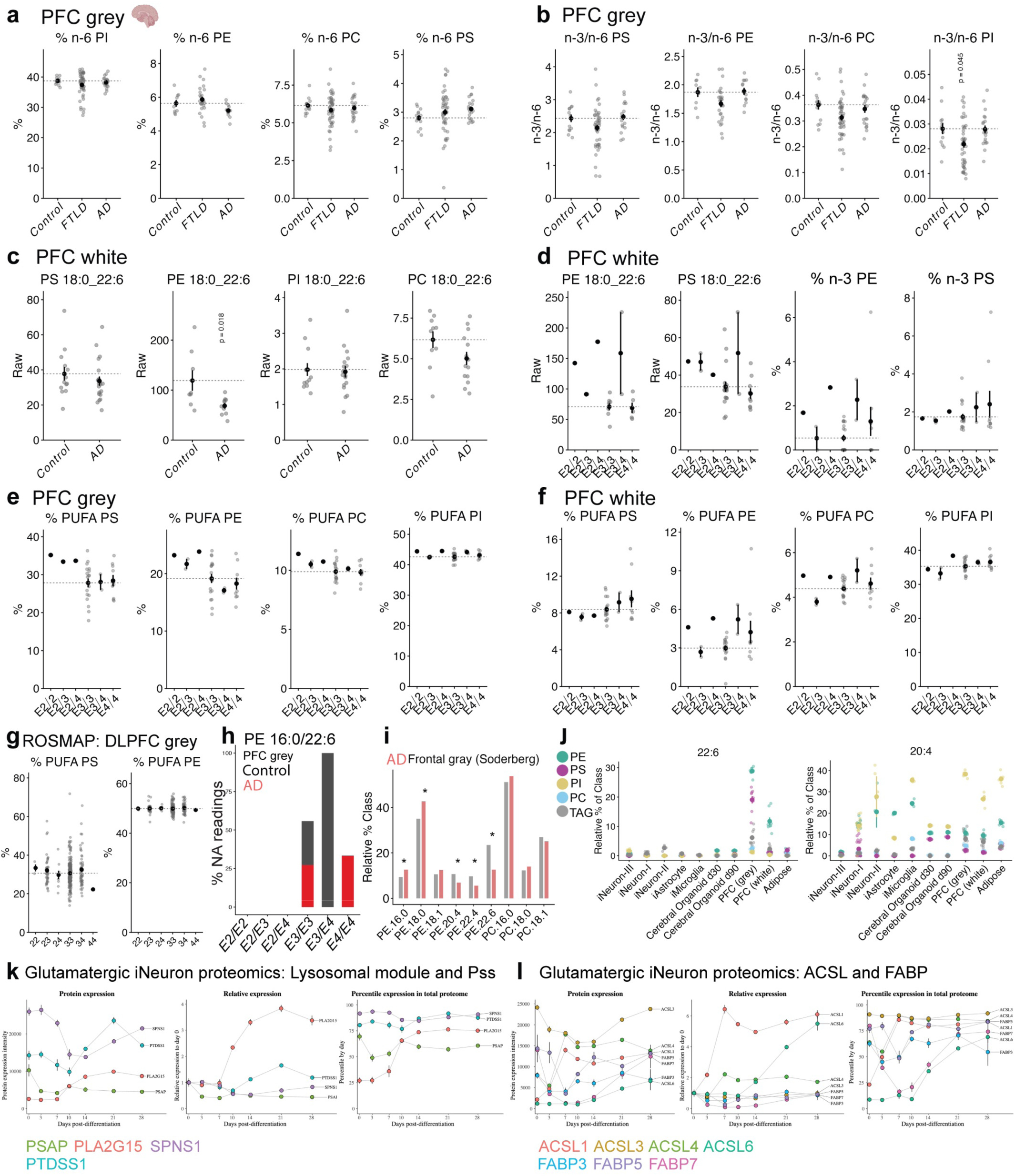
FTLD and AD disease alterations of brain PUFAs. **a,** Neurolipid Atlas prefrontal cortex (PFC) grey matter from controls, frontotemporal lobar dementia (FTLD), and Alzheimer’s patients (AD) analyzed for % n-6 ratios in PS, PE, PI, and PC. **b,** PFC grey matter n-3/n-6 ratios for PS, PE, PC, and PI from the Neurolipid atlas. **c,** PFC white matter raw 18:0_22:6 values from Neurolipid atlas (only AD patients had white matter profiled). p-values represent BH-corrected two-sided t-tests against control. **d,** PFC white matter 18:0_22:6 and % n-3 for PE and PS lipids, stratified based on APOE status. **e,** PFC grey matter % PUFA for PS, PE, PC, and PI, stratified by APOE status. **f,** PFC white matter % PUFA for PS, PE, PC, and PI, stratified by APOE status. **g,** ROSMAP data^94^ for %PUFA PS and % PUFA PE, stratified by APOE status. **h,** proportion of 0/NA values for PE 16:0/22:6 in the Neurolipid atlas (red = AD, black = control). **i,** data replotted from tables in^95^, with control and AD (red) relative % fatty acid chain composition across frontal grey matter. **j,** relative % contribution of essential fatty acids 22:6 or 20:4 to PS (magenta), PE (green), PC (blue), PI (gold), or TAG (grey) from the same dataset as Fig. 8h-i. n = 8-10 for Soderberg data; for Neurolipid atlas, n = 13 controls, 46 FTLD, and 20 AD patients for grey matter. For APOE status in the Neurolipid atlas: E2/E2 n=1; E2/E3 n =2; E2/E4 n =1; E3/E3 n =18; E3/E4 n =2; and E4/E4 n =9. For ROSMAP APOE status: E2/E2 n = 3; E2/E3 = 42; E2/E4 = 9; E3/E3 = 202; E3/E4 = 59; E4/E4 = 1. **k,** reanalysis of iNeuron longitudinal proteomics^101^ for PSAP, PLA2G15, SPNS1, and PTDSS1. **k,** reanalysis of iNeuron longitudinal proteomics^101^ for ACSL1, ACSL3, ACSL4, ACSL6, FABP3, FABP5, and FABP7. Data are represented by mean and SEM.

## Contributions

Conceptualization: RJL, IMRS, JPV. Methodology: RJL, IMRS, EKT, CIT, AD, MTC, JPV. Investigation: RJL, IMRS, EKT, AB, EBH, PC, ZZ, JSCR, JCF, TTP, CIT, JPV. Visualization: RJL, JPV. Funding acquisition: AD, VM, JPV. Project administration: VM, JPV. Supervision: JPV. Writing: RJL, JPV.

## Acknowledgments

Research done at UCSF is conducted on Ramaytush Ohlone land. We thank Vaughen lab members for discussion, Michael Hoch, Sean Sweeny, Liqun Luo, Jairaj Acharya, Quan Yuan, Kate O’Connor Giles, Taylor Jay, and Chih-Chiang Chan for sharing flies; Sebastian Sorge for HolFast discussion; Chih-Yu Chen for sharing raw ROSMAP data; Yerbol Kurmangaliyev for scRNAseq discussion; Martin Kampmann for iNeuron discussion; Saul Villeda for qRT-PCR access; and Rushika Perera, Leanne Jones, and Andrew Yang for confocal access. This work was supported by the Sandler PBBR program (JPV), the Simons Foundation (SFI-AN-NC-AB-Research-00018030 to JPV), the American Parkinson Disease Association (JPV), the Shurl and Kay Curci Foundation (EBH), the NSF GRFP (JSCR), NIH T32-HD007470 (TTP), NIH R21 NS144615 (JPV), and by the Electron Microscopy Core Facility at CU Anschutz (1S10OD036258-01). ZZ was supported in part by Simons Foundation (SFI-AN-Director-00017753 to Olga Troyanska). We gratefully acknowledge Flybase^169,170^, Bloomington Drosophila Stock Center (NIH P40OD018537), and the Harvard TRiP Center (R24 OD030002)^171^.

